# Pyruvate promotes ciliogenesis bypassing *IFT88* dependency and attenuates DSS-induced colitis

**DOI:** 10.64898/2025.12.17.694572

**Authors:** Maya Sarieddine, Ilaria Cicalini, Damiana Pieragostino, Federica Dimarco, Matthieu Lacroix, Krzysztof Rogowski, Valérie Pinet, Michael Hahne

## Abstract

Primary cilia (PC) are important signaling rheostats, yet their biology in colon remains understudied. We previously reported that the presence of PC on colonic fibroblasts in mice (CF) modulates their susceptibility to colitis. Here, we demonstrate that extracellular pyruvate levels influence both ciliary length and ciliogenesis in CF. Pyruvate supplementation to CF enhanced tubulin and histone acetylation, with the latter promoting MAPK signaling and tubulin detyrosination within PC. MAPK-inhibition reduced tubulin detyrosination and shortened ciliary length, while inhibition of α-tubulin acetylation specifically affected ciliogenesis. *Col6a1cre-Ift88flx/flx* mice, lacking ciliary *Ift88* gene in *Col6a1*-expressing CF, displayed reduced ciliogenesis and increased susceptibility to DSS-induced colitis. Surprisingly, in primary CF cultures from these mice, pyruvate supplementation restored PC formation. Moreover, pyruvate administration via drinking water rescued PC formation in *Col6a1cre-Ift88flx/flx* mice and attenuated DSS-induced colitis. These findings identify pyruvate as a regulator of PC biology in CF and as a therapeutically relevant factor in colitis.

**Graphical Abstract:** Pyruvate promotes ciliogenesis bypassing *IFT88* dependency and attenuates DSS-induced colitis

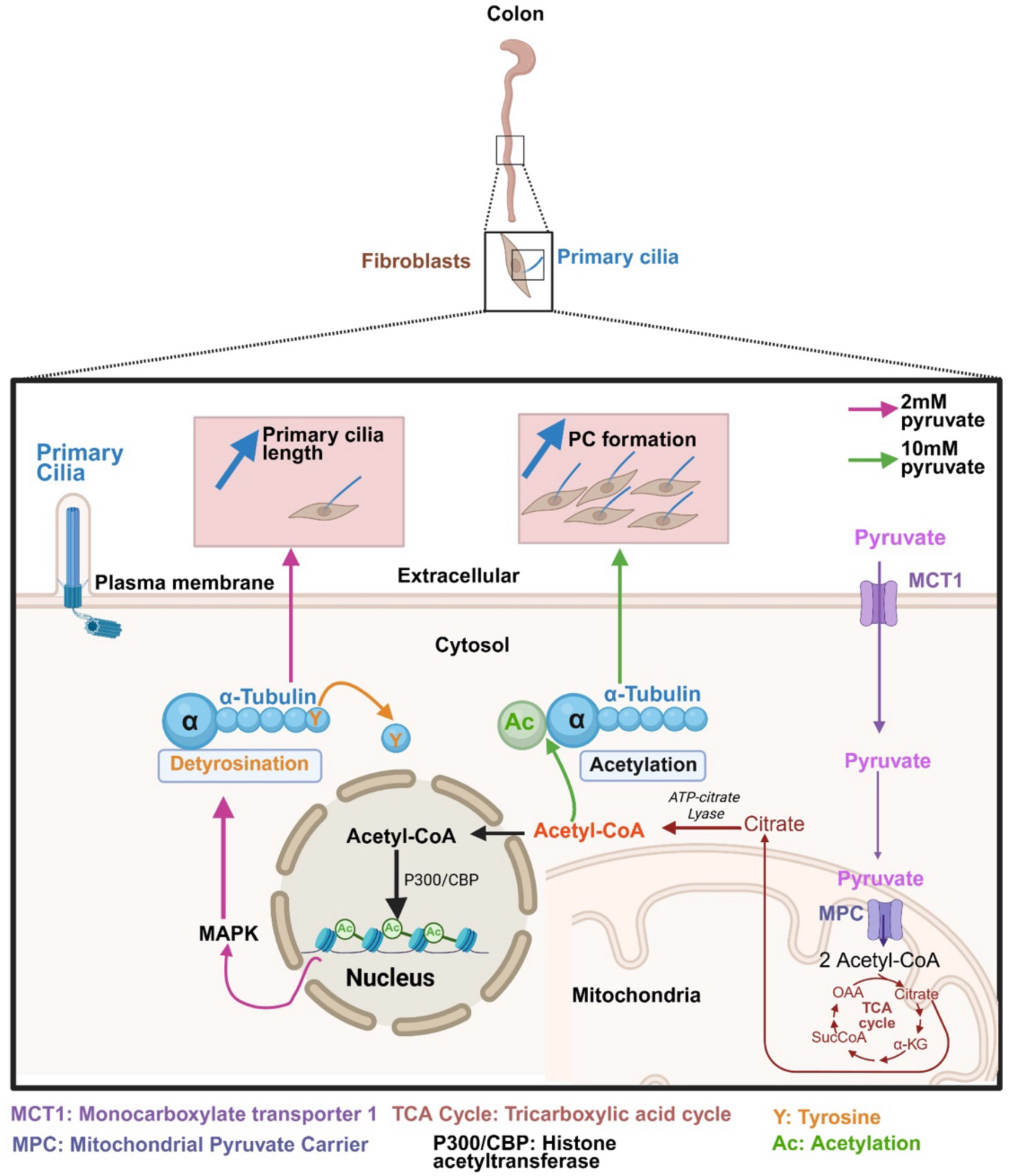

## Introduction

The crosstalk between epithelium lining the crypts and neighboring fibroblasts is crucial for maintaining intestinal homeostasis and the regulation of pathological responses. An underexplored player in this signaling interplay is the primary cilium, an antennae-like structure protruding from the surface of many mammalian cells that serves as a signaling sensor. The axoneme, the core of the primary cilium, is composed of nine microtubule doublets assembled from α- and β-tubulin heterodimers in a polarized manner, forming a cylindrical scaffold. This structure is subject to a broad variety of tubulin post-translational modifications (PTMs), such as acetylation, detyrosination, glutamylation and glycylation ^1^. These PTMs are essential regulators of PC stability and function, and a concept emerged that the interplay between different PTMs dictates the functional output of PC in mammals ^2,3^.

PC have been extensively studied in tissues where they are prominent and functionally critical, particularly in the kidney and brain ^4,5^. We previously identified a role for PC in the colon biology ^6^. Notably, we found a reduced numbers of PC on colonic fibroblasts (CF) during acute colitis as compared to those of untreated mice. These data revealed a link between the presence of PC and colonic inflammation. Importantly, our observations in the murine model of colitis were confirmed by data obtained with inflamed regions of biopsies derived from patients with ulcerative colitis (UC), which displayed a reduced number of PC in comparison to the neighboring normal tissue.

To assess whether a reduction of PC numbers in the colon increases the susceptibility to induced colitis, we employed a reference models for PC loss. Specifically, we used mice carrying floxed allele of the intraflagellar transport protein 88 (*Ift88^flx/flx^*), that has been reported to be absolutely required for PC assembly and function ^7,8^. This mouse strain was crossed with collagen6a1(Col6a1)cre mice to direct gene deletion in Col6a1-expressing CF ^6^. Colons of *Col6a1cre-Ift88^flx/flx^*mice displayed decreased number of PC and increased susceptibility to chemically induced acute colitis underpinning the link between PC presence and predisposition to intestinal inflammation.

Length, number, and function of PC are dynamically regulated by both, environmental and intracellular cues. Among these, nutrients, soluble factors and ions are important modulators of ciliogenesis. For instance, a study in human telomerase-immortalized retinal pigment epithelial cells (RPE1), a frequently used cell model for ciliary research, reported that elevated extracellular calcium shortens PC length by depolymerizing axonemal microtubules ^9^. On the other hand, glucose deprivation promotes ciliogenesis in RPE1 cells ^10^. More recently, Steidl *et al.* demonstrated that glutamine depletion, but not glucose deprivation, promotes PC elongation in mouse embryonic fibroblasts (MEFs) and immortalized mouse inner medullary collecting duct (IMCD3) cells ^11^. Collectively, these findings indicate that extracellular factors modulate PC biology in a cell type–specific manner.

Here, we investigated how environmental factors might influence PC biology in the colon. Our study identified pyruvate as an important regulator of ciliogenesis and PC length in CF.

## Results

### Sodium pyruvate promotes PC elongation and ciliogenesis in serum starved colonic fibroblasts

Serum starvation is a commonly used method to induce PC formation in cultured cells. We have previously established *ex vivo* primary murine colonic fibroblast (CF) cultures that allow to explore PC biology at low passage numbers ^6^. When subjected to serum starvation in RPMI medium, these cultures accurately recapitulate the proportion of PC-positive fibroblasts observed in the colon of wild-type mice ^6^. Since serum starvation is known to promote primary cilium elongation in multiple cell types, including mouse embryonic fibroblasts ^11^, we quantified primary cilium length in serum-starved CFs. As shown in Fig. 1A, PC length was similar in non-starved and serum-starved colonic fibroblasts, when performed in RPMI medium.

**Figure 1:**
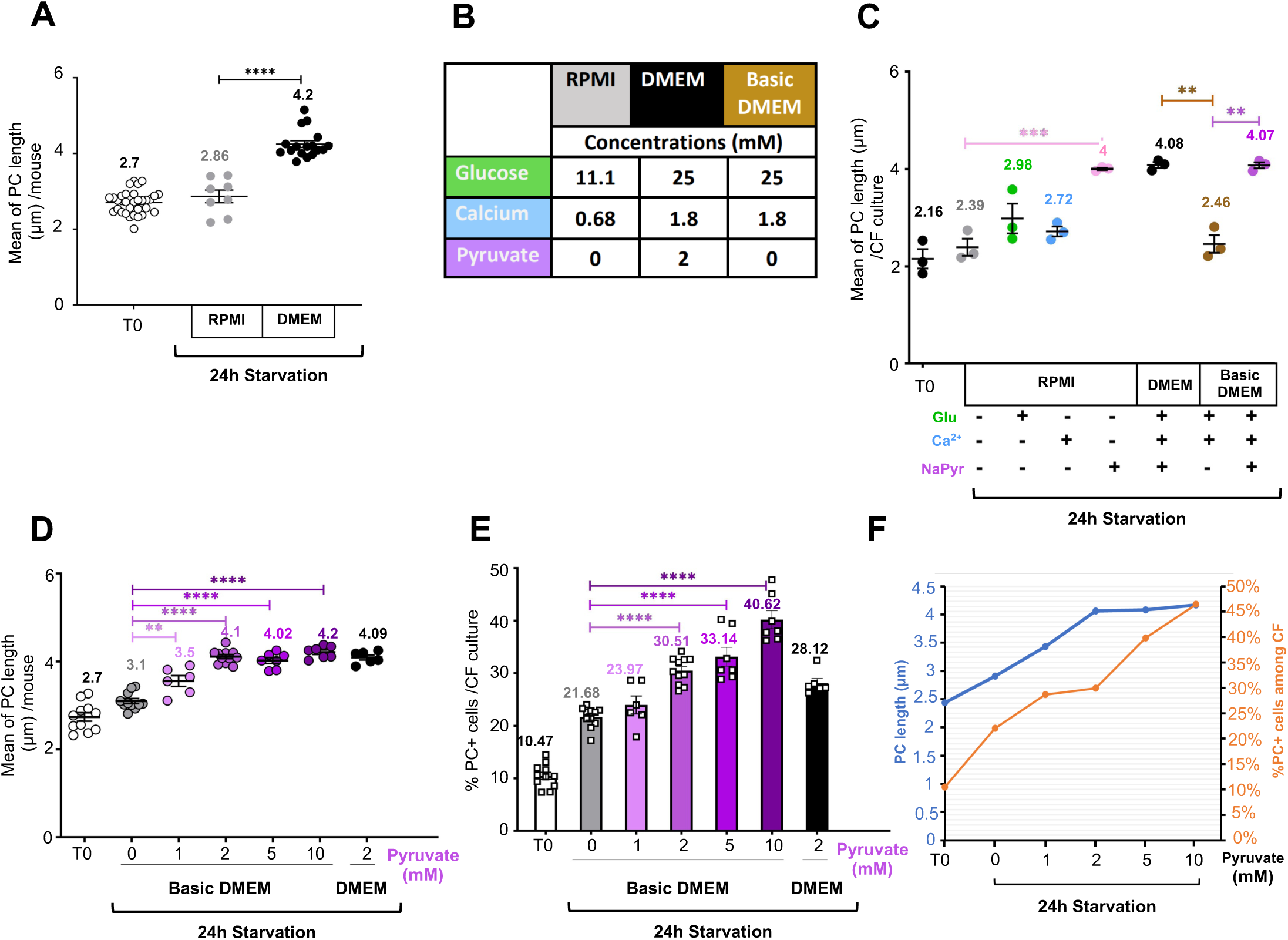
Pyruvate regulates PC length and ciliogenesis. **(A)** Quantification of PC length per colonic fibroblasts after 24h starvation, in both RPMI and DMEM medium. Each point shows the mean cilium length of one CF culture obtained from an individual mouse. **(B)** Comparison of component concentrations in RPMI and DMEM media. **(C)** Quantification of PC length per CF culture after 24h starvation in RPMI, RPMI complemented with either Glucose (Glu, 0.14mM), Ca^2+^ ions (0.93 mM), or sodium pyruvate (2mM), or in complete or basic DMEM. **(D-F)** Impact of pyruvate concentrations (0, 1, 2, 5, 10 mM) on PC length (**D)**, and number of PC-positive cells (**E)**. Overlay graph of pyruvate concentration effects on PC length (blue) and PC⁺ cell number (orange) **(F)**. Statistical analysis in **(A, C, D, E)**: **p < 0.01, ***p < 0.001, ****p < 10^−4^ by two-tailed unpaired t-test.

Since PC length is regulated by nutrients and extracellular factors, we tested whether different media compositions could modulate PC length. Replacing RPMI with the nutrient-richer DMEM medium led to the formation of significantly longer cilia (Fig. 1A). Compared with RPMI, DMEM contains higher levels of glucose, Ca²⁺ ions, and pyruvate (Fig. 1B). To test which of these components accounted for the increased PC length, we serum starved CFs either in RPMI alone or in RPMI supplemented with either glucose, calcium chloride (CaCl_2_) or pyruvate at the concentrations present in DMEM. Addition of either glucose or calcium chloride had no significant effect on PC length (Fig. 1C and Supplementary Fig. 1). Strikingly, the addition of sodium pyruvate elongated PC to an extend comparable to that observed with DMEM. Based on these findings, all subsequent experiments were carried out in DMEM containing glucose and Ca²⁺ but lacking pyruvate (hereafter referred to as basic DMEM), unless otherwise specified. Indeed, the ciliary length in the cells starved in basic DMEM supplemented with 2mM pyruvate was comparable to the one observed in full DMEM (Fig. 1C).

Additional testing using basic DMEM supplemented with increasing concentrations of sodium pyruvate (1, 2, 5, and 10 mM) showed that PC reached a maximum length of 4 µm at 2 mM, with higher pyruvate concentrations having no further effect on ciliary length (Fig. 1D). Next, we tested whether sodium pyruvate levels could also regulate the number of PC-proficient CF during serum starvation. We observed, a dose-dependent increase in the number of ciliated CFs (Fig. 1E), showing that sodium pyruvate apart from regulating ciliary length also affects the levels of ciliation. In contrast, the addition of glucose or Ca^2+^ ions had no effect on the number of ciliated cells (Supplementary Fig. 2).

### Pyruvate modulates PC occurrence through tubulin acetylation

Pyruvate is a major source of acetyl-CoA, which is the primary donor for tubulin acetylation catalyzed by α-tubulin N-acetyltransferase 1 (ATAT1) (Fig. 2A) ^12^. Therefore, we tested whether the concentration of sodium pyruvate could modulate the acetylation status of PC in starved CFs. We observed a dose-dependent increase in ciliary acetylation, which was correlated with increasing concentrations of sodium pyruvate during starvation (Fig. 2B). Since, acetyl-CoA is generated from pyruvate in mitochondria, we investigated pyruvate transport using two inhibitors, AZD3965 and UK5099. AZD3965 is a selective inhibitor of MCT1, which transports pyruvate across the plasma membrane and does not directly affect mitochondrial transport of pyruvate. Conversely, UK5099 is a specific inhibitor of the MPC localized at the inner membrane of mitochondria. Addition of either of the two inhibitors, reduced the length of PC to that observed in serum-starved CF in the absence of pyruvate (Fig. 2C).

**Figure 2:**
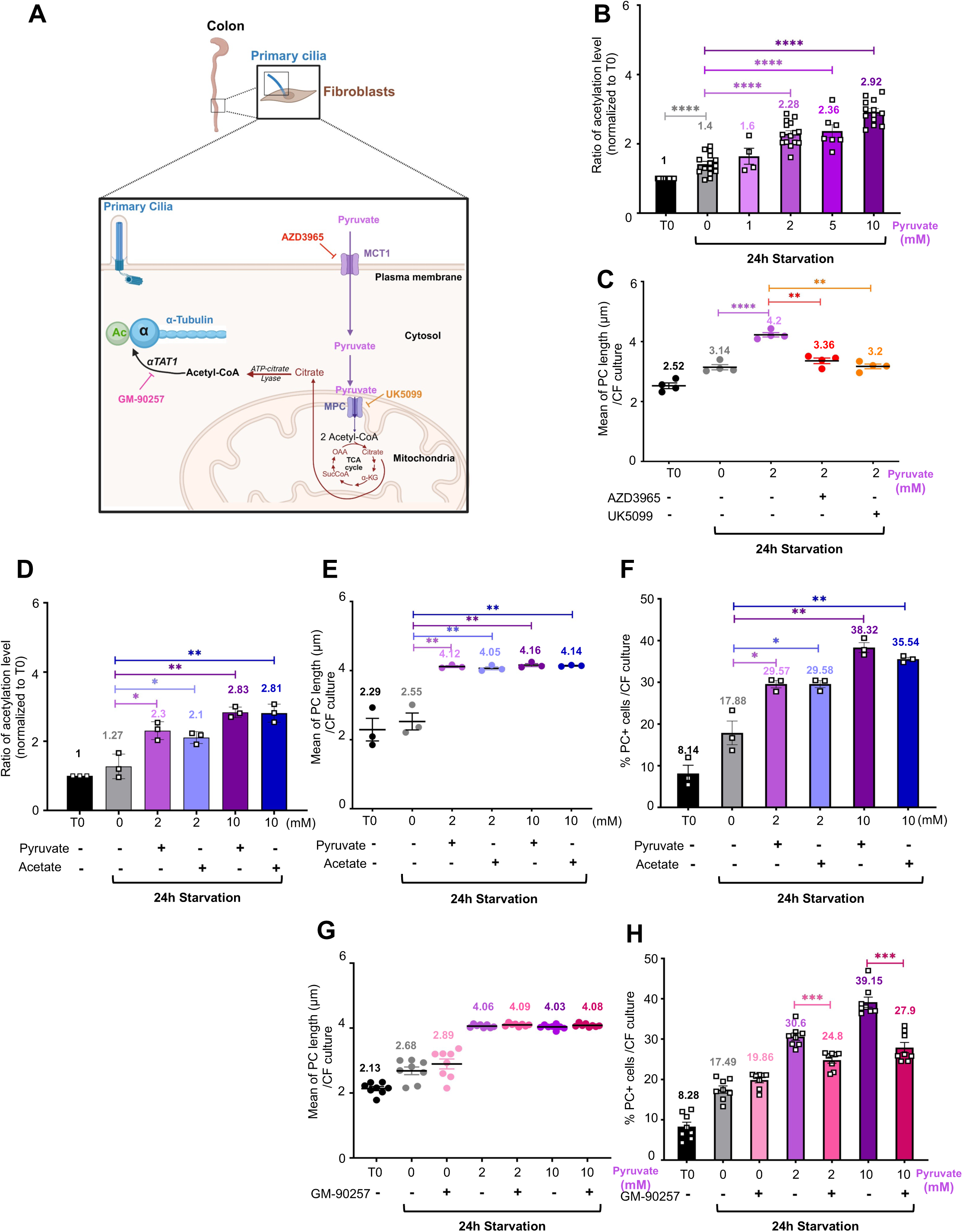
Pyruvate regulates PC presence by modulating tubulin acetylation. **(A)** Schematic illustration how pyruvate can promote tubulin acetylation. Extracellular pyruvate is transported across the plasma membrane via the monocarboxylate transporter (MCT1), then entering mitochondria through the Mitochondrial Pyruvate Carrier (MPC). In the mitochondria, pyruvate is converted to acetyl CoA which fuels the TCA cycle, citrate exits the mitochondria and is converted back to cytosolic acetyl-CoA, which serves as the acetyl donor for α-tubulin acetyltransferase (αTAT1) to acetylate α-tubulin in primary cilia microtubules. Pharmacological inhibitors targeting this mechanism are indicated: AZD3965 targets MCT1, UK5099 blocks MPC, and GM-90257 inhibits ATAT1. **(B-H)** CF were serum starved in basic DMEM supplemented with indicated concentrations of Pyruvate **(B)** Impact of pyruvate increasing concentrations (0, 1, 2, 5, 10 mM) on PC acetylation. The acetylation level was quantified using integrated density/area measurements in ImageJ. The area was delineated using Arl13b staining, and the density of acetyl tubulin staining was quantified in this area **(C)** Impact of pyruvate transport inhibitors AZD3965 (200nM) and UK5099 (50µM) on PC length. **(D-F)** Impact of sodium pyruvate or sodium acetate supplementation on PC acetylation (**D)**, PC length **(E)** and number of PC-positive CF (**F**). **(G-H)** Effect of acetylation blocking using the inhibitor GM-90257 (500nM) on both PC length (**G)** and number of PC-positive CF (**H)**. Statistical analysis **(B-H)** : two-tailed unpaired t-test; *p<0.05, **p< 0.01, ***p< 0.001, ****p<0.0001.

Another precursor for acetyl-CoA in experimental settings is sodium acetate, which enters the cells mainly through passive diffusion. Indeed, addition of 2mM sodium acetate during serum starvation elevated acetylation levels in PC (Fig. 2D) and increased PC length (Fig. 2E) as well as number of PC-positive CFs (Fig. 2F) similarly to the levels observed in the presence of 2mM sodium pyruvate. These data provide further evidence that the increase in ciliary length and number is most likely mediated by acetyl-CoA. Finally, we treated the cells with GM-90257, an inhibitor of ATAT1, which catalyzes acetylation of α-tubulin on lysine 40 (Fig. 2A). GM-90257 treatment had no effect on pyruvate-triggered PC elongation (Fig. 2G), however it reduced the number of PC containing CFs (Fig. 2H). In summary, pyruvate appears to regulate PC number in CF through tubulin acetylation, but not the PC length.

### Pyruvate induces elongation of PC through the activation of the MAPK pathway

To gain insights in the underlying mechanisms of pyruvate-triggered PC elongation in CF cultures, we performed RNA-seq analysis of cells grown in the absence or presence of 2mM sodium pyruvate. Supervised analysis revealed 1452 genes differentially regulated in CF cultured in the presence of 2mM pyruvate as compared to control condition, among which 1251 genes were upregulated and 201 genes were downregulated, using a p-value ≤ 0.05 and a Log2 fold change ≥ 1 as threshold (Fig. 3A). The gene expression signature was then explored by GSEA taking Kyoto Encyclopedia of Genes and Genomes (KEGG) as data source, which found Mitogen-Activated Protein Kinase (MAPK) signaling pathway to be the most upregulated (Fig. 3B). We found 34 genes of the MAPK pathway among the leading-edge subset enriched in our analysis (Fig. 3C). In addition, we performed proteomics analysis of fibroblasts starved in absence or presence of 2mM pyruvate, which also indicated an upregulation of MAPK signaling. More precisely, we found the Rapidly Accelerated Fibrosarcoma (RAF)/MAPK cascade with ERK1 among the most upregulated proteins (Supplementary Table 1). Furthermore, following pyruvate treatment, we also observed an upregulated PC signature by mass spectrometry (Supplementary Table 1). These data were corroborated by the aforementioned RNA sequencing experiment, which displayed increased expression of genes critically involved in primary cilia biology such as BBSome components, hedgehog signaling pathway genes, and additional PC-associated genes (Supplementary Fig. 3).

**Figure 3:**
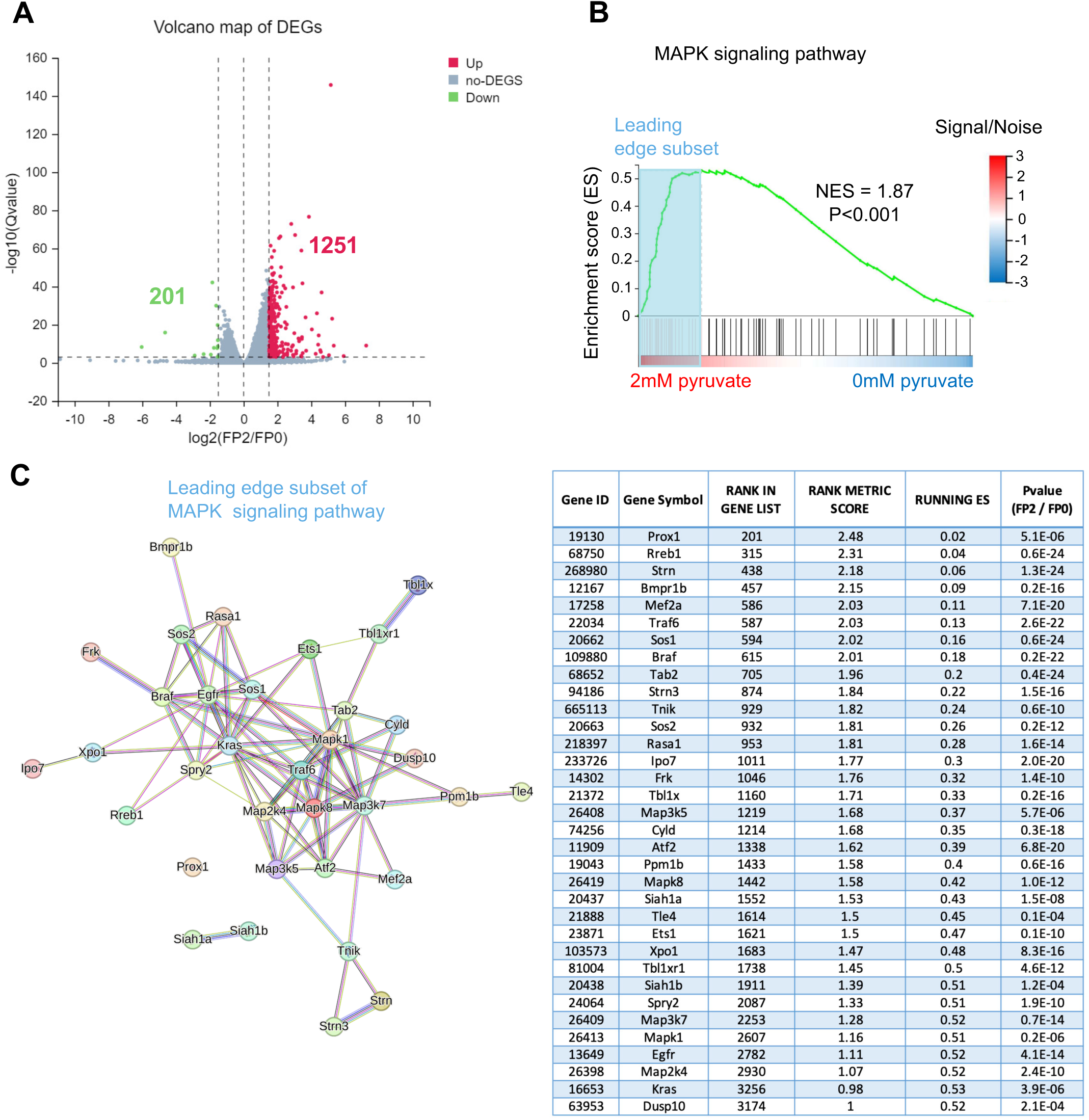
Enrichment of MAPK signaling pathway in serum-starved colonic fibroblasts in the presence of 2mM pyruvate. **(A)** Volcano plot showing 1452 differentially expressed genes (DEG) in 24h serum starved CF cultures (n=3) in the absence of presence of 2 mM pyruvate identified by RNAseq. 1251 upregulated and 201 downregulated genes are shown in red and green, respectively, displaying a q-value ≤ 0.05 and Log2 fold change ≥ 1. **(B)** Gene Set Enrichment Analysis (GSEA) plot of MAPK signaling displaying the enrichment score (ES) presented by the green line across the ranked gene list with the leading-edge subset indicated by blue rectangle representing genes contributing most to the enrichment score, including the normalized enrichment score (NES) and adjusted p-value provided. **(C)** Search Tool for the Retrieval of Interacting Genes (STRING) network (left) and table (right) illustrating the composition of the leading-edge subset of the MAPK signaling pathway, highlighting core genes driving this enrichment.

To validate the implication of the MAPK pathway in pyruvate-triggered PC biology we used PD98059, an inhibitor of the MAP kinase cascade blocking the activation of the kinases MEK1/2. The addition of PD98059 reduced pyruvate-triggered PC elongation (Fig. 4A), but not the number of PC-positive CF (Fig. 4B). To exclude the off-target effect of PD98059, we used an additional inhibitor of MAPK signaling, i.e., SCH772984, which blocks ERK1/2. Consistently with the effects of PD98059, the inhibition of ERK1/2 reduced pyruvate-related increase in PC length, but had no effect on the number of ciliated CFs (Fig. 4C and 4D). Finally, the inhibition of the MAPK pathway regardless of the inhibitor used, had no significant effect on PC acetylation (Supplementary Fig. 4).

**Figure 4:**
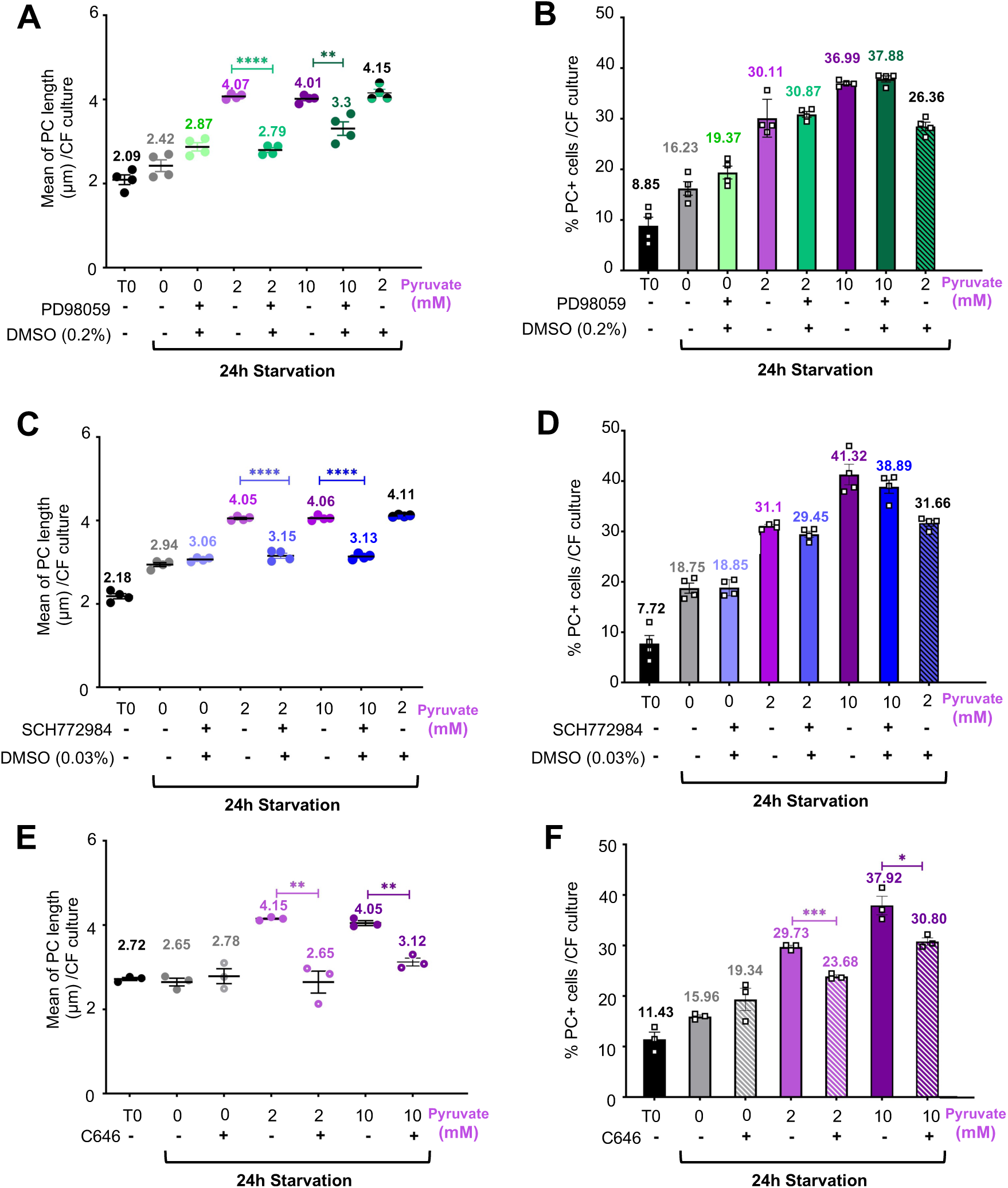
MAPK modulates PC length while histone acetylation affects both PC length and ciliogenesis in serum starved CF. PC length **(A, C, E)** and number of PC-positive cells **(B, D, F)** in 24h serum starved CF cultures in absence or presence of 2mM sodium pyruvate and supplemented with indicated concentrations of MEK1/2 inhibitor PD9805 **(A, B),** ERK1/2 inhibitor SCH772984 (**C, D)** and histone acetylation EP300 inhibitor C646 (**E, F).** Statistical analysis: *p<0.05, **p< 0.01, ***p< 0.001, ****p<0.0001 by two-tailed unpaired t-test.

In addition to functioning as a co-factor for tubulin acetylation, acetyl-CoA also supports histone acetylation, which can regulate gene transcription ^13^. This prompted us to test whether changes in histone acetylation could be at the origin of the pyruvate-induced increase in the transcript levels of MAPK pathway components. Consistent with this model, the GSEA of pyruvate-treated CFs indicated elevated histone acetylation activity with a prominent upregulation of the acetyltransferases EP300, Kat6 and Jade1 (Supplementary Fig. 5). Accordingly, the addition of the EP300 inhibitor C646 to serum starved CF reduced pyruvate-triggered PC length (Fig. 4E), as well as number of PC-proficient CFs (Fig. 4F). A similar outcome was observed with the EP300 inhibitor A-485, which reduced both PC length and the percentage of PC-proficient cells (Supplementary Fig. 6). Interestingly, inhibition of the histone acetyltransferase EP300 reduced pyruvate-induced PC elongation to a degree comparable to MAPK inhibition and also decreased the number of pyruvate-responsive ciliated CFs, resembling the effects observed upon ATAT1 inhibition. These results suggest that EP300 functions upstream of, or in concert with, the MAPK pathway and ATAT1 in the regulation of pyruvate-dependent ciliary assembly and elongation.

### Pyruvate modulates tubulin detyrosination status

Another prominent tubulin modification, which accumulates on ciliary microtubules is detyrosination ^14^. Thus, we investigated whether this modification could play a role in pyruvate induced alterations of PC biology in CF during serum starvation. We observed that detyrosination levels were elevated in the PC of serum-starved CF cells with 2 mM pyruvate, but no further enhancement was observed at 10 mM (Fig. 5A). To test whether pyruvate facilitates crosstalk between tubulin acetylation and detyrosination in PC, we treated cells with the ATAT1 inhibitor GM-90257. We found that GM-90257 had no effect on pyruvate-induced increase in detyrosination of PC-positive CFs under starvation conditions, regardless of pyruvate concentration (Fig. 5B).

**Figure 5:**
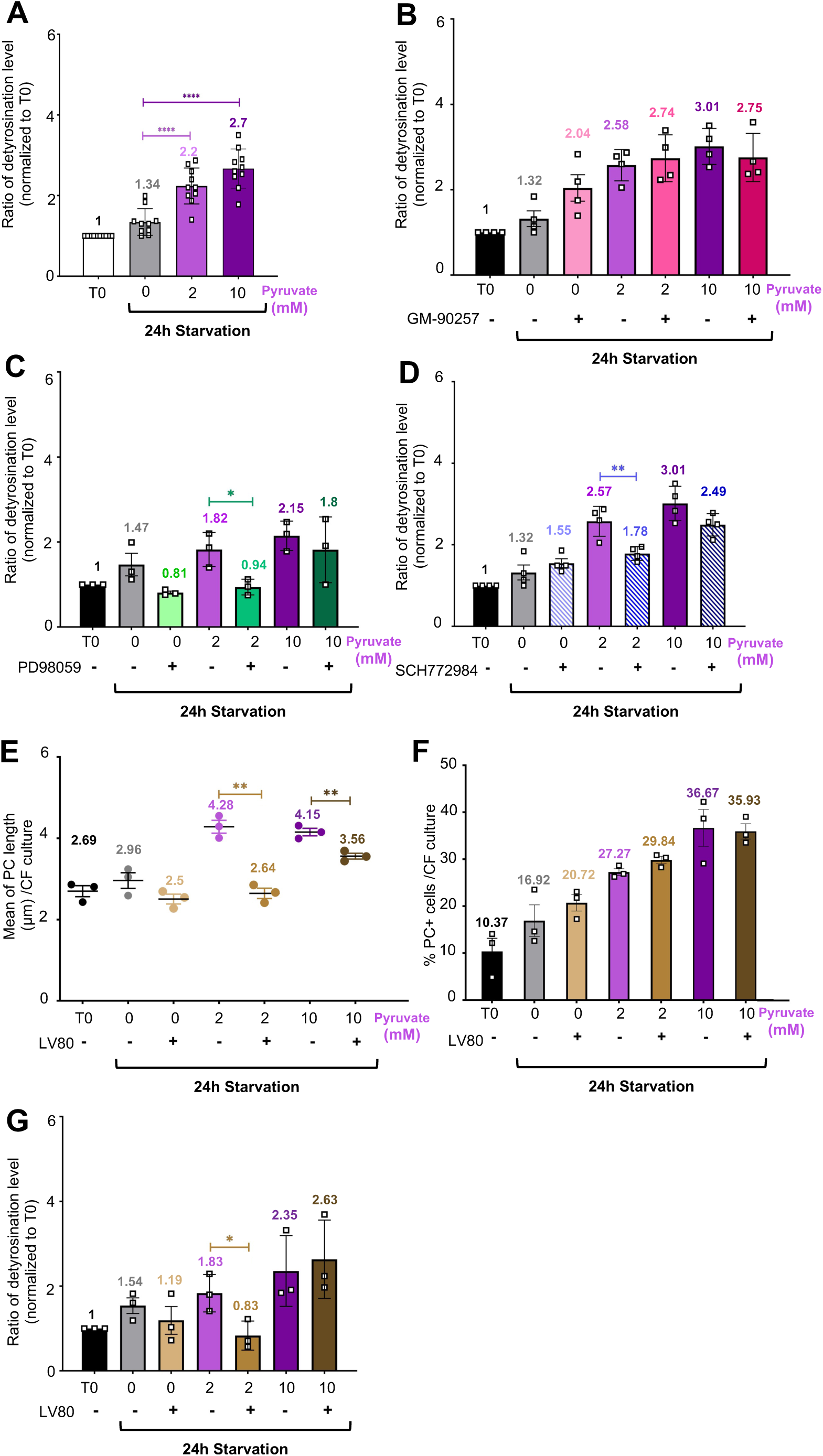
Detyrosination of tubulin is implicated in the regulation of PC length in serum starved CF. **(A)** In presence of pyruvate during CF serum-starvation, PC detyrosination increased in a dose dependent manner. **(B-D)** Graphs showing the impact of acetylation inhibition using GM-90257 **(B)** and MAPK inhibition using inhibitors PD98059 **(C)** and SCH772984 **(D)** on the detyrosination at PC level. **(E-F)** Graphs showing the impact of the detyrosination inhibitor LV80 on PC length **(E)**, number of PC-positive cells **(F)** and detyrosination level of PC **(G)** in 24h serum starved colonic fibroblasts cultures. **(A, C, D, E, G)** Statistical analysis: *p<0.05, **p< 0.01, ****p<0.0001 by two-tailed unpaired t-test.

Since pyruvate induces elongation of PC at 2 mM through activation of the MAPK pathway, we next assessed whether MAPK signaling influences detyrosination levels. Using the MAPK and ERK1/2 inhibitors PD98059 and SCH772984, respectively, we found that both inhibitors reduced detyrosination levels of PC in serum-starved CFs cultured with 2 mM pyruvate (Fig. 5C and Fig. 5D), but had no effect at 10 mM pyruvate. These findings suggest that MAPK promotes detyrosination and thus PC length.

The main class of enzymes, which catalyze detyrosination are vasohibins (VASH1 and VASH2)^15,16^. Therefore, we used a specific VASH inhibitor, LV80 to assess the role of detyrosination in the regulation of ciliary length in serum starved CFs. We found that LV80-mediated VASH inhibition reduced PC length but did not affect the number of ciliated cells (Fig. 5E and 5F). Furthermore, while the addition of LV80 efficiently reduced PC detyrosination at 2 mM pyruvate, it had no effect in the cells grown at 10 mM (Fig. 5G). Altogether, these results suggest that VASH-mediated detyrosination is specifically involved in the regulation of PC length.

### Pyruvate can overcome IFT88-dependency of PC

We previously described that colon of *Col6a1cre-Ift88^flx/flx^* mice display a lower number of PC-positive CFs in comparison with colon of control animals ^6^. In addition, we demonstrated that CFs cultures grown in serum-free RPMI medium recapitulate the different number of PC in CF observed in colons of control and *Col6a1cre-Ift88^flx/flx^*mice ^6^. These data were in agreement with the reported essential role of *Ift88* in PC assembly and maintenance ^17^.To monitor collagen6a1 expressing cells we crossed *Col6a1cre-Ift88^flx/flx^* mice with *ROSA^mT/mG^*reporter mice to introduce membrane-targeted GFP in cre-expressing cells (Fig. 6A). Immunofluorescence analysis showed a gradient of Col6a1 expression in the lamina propria of the colon (Supplementary Fig. 7). As expected from our previous data, CF isolated from *Col6a1cre-Ift88^flx/flx^-ROSA^mT/mG^* mice displayed reduced number of PC as compared to those isolated from control *Ift88^flx/flx^-ROSA^mT/mG^*mice when serum starved in RPMI medium (Fig. 6B). However, when *Ift88*-proficient and -deficient CFs were serum starved in DMEM (i.e. in the presence of 2mM pyruvate) the number of cilia in both cell types was comparable (Fig. 6B). We next validated IFT88 deficiency in CF of *Col6a1cre-Ift88^flx/flx^-ROSA^mT/mG^*mice. Accordingly, immunostaining revealed the presence IFT88 protein in PC of wild-type CFs, whereas GFP-positive, i.e. collogen6a1 expressing CF derived from *Col6a1cre-Ift88^flx/flx^-ROSA^mT/mG^* mice were negative for *IFT88* (Fig. 6C). Taken together these results show that the addition of pyruvate rescues the IFT88 deficiency in CFs. To assess whether pyruvate could also restore PC presence in the colon of *Ift88*-deficient mice, we added for 8 weeks 0.2M sodium pyruvate or NaCl in the drinking water of newly weaned *Col6a1cre-Ift88^flx/flx^-ROSA^mT/mG^*and control mice (i.e. at 3 weeks of age). Addition of sodium pyruvate or osmolarity matched NaCl did not induce any apparent physiological changes in the mice as shown by the body weight curves of the animals (Supplementary Fig. 8). *Ift88*-deficient mice are smaller than control mice at birth. Indeed, *Ift88* was shown to be essential for development ^17^, which might explain the delayed growth in *Ift88*-deficient mice at early age. However, *Ift88*-deficient mice compensated for the reduced weight in comparison to controls within one-week post-weaning (Supplementary Fig. 8). At the end of the protocol, mice were sacrificed, and their colons analyzed for presence of PC by immunofluorescence (Fig. 6D). In agreement with our previous report ^6^, *Ift88*-deficient animals displayed a significantly lower number of colonic PC-positive fibroblasts as compared to controls, when 0.2 M NaCl was added in drinking water. Strikingly, the addition of pyruvate in the drinking water restored the percentage of PC-positive fibroblasts in *Col6a1cre-Ift88^flx/flx^-ROSA^mT/mG^* mice to similar levels as found in control animals (Fig. 6E). Furthermore, the control mice treated with pyruvate exhibited a slight increase of PC-positive fibroblast number (Fig. 6E) accompanied by a significant extension of PC length (Fig. 6F). Thus, our data show that pyruvate treatment not only compensates for *Ift88*-deficiency by increasing the number of colonic PC-positive fibroblasts *in vitro* as well as *in vivo,* but also increases ciliary length in control mice recapitulating the effect observed *in vitro*.

**Figure 6:**
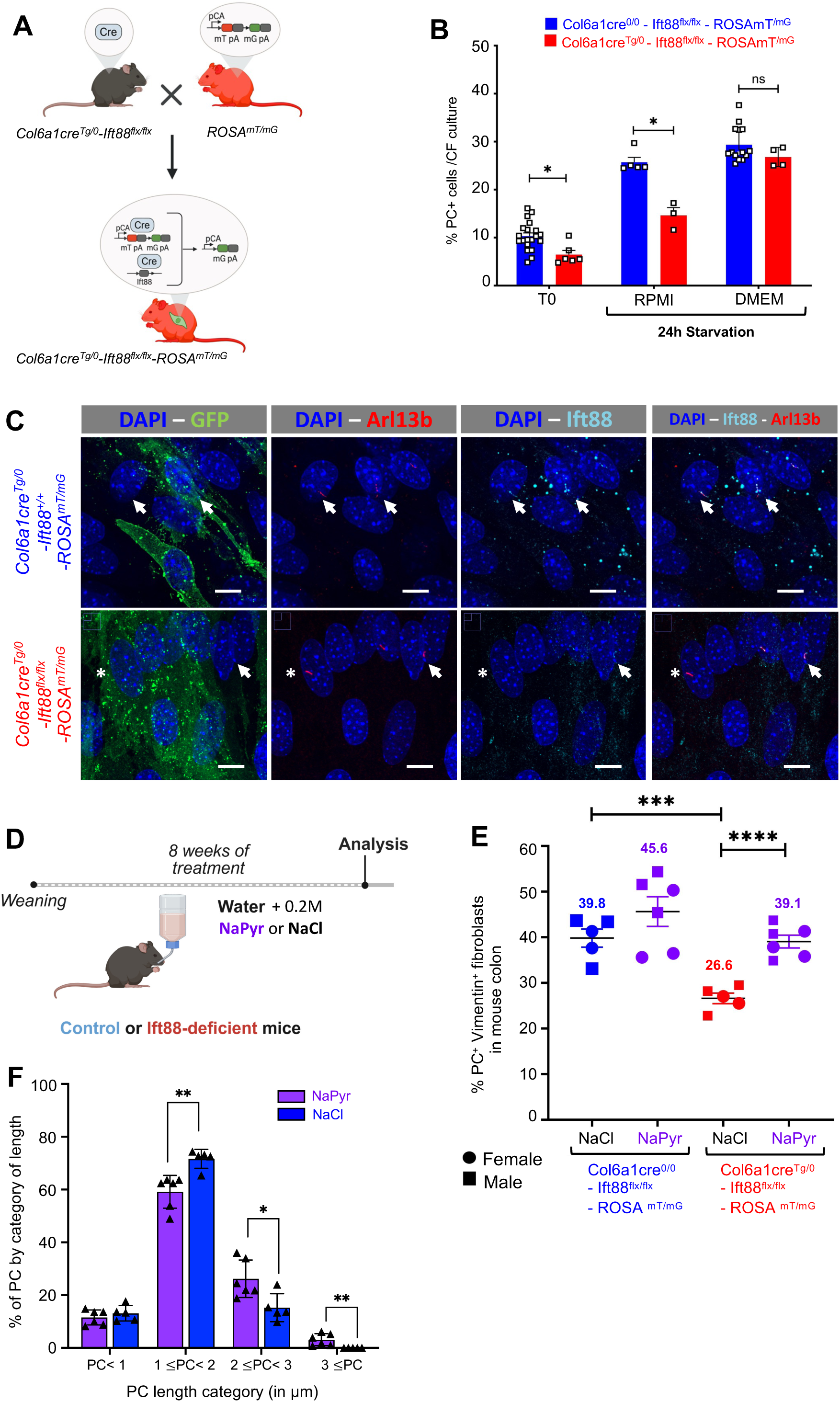
Pyruvate can overcome Ift88 dependency of primary cilia. **(A)** Schematic illustration of the breeding strategy employed to generate *Col6a1cre-Ift88^flx/flx^-Rosa ^mT/mG^* mice deleting *Ift88* specifically in collagen 6a1-positive colon fibroblasts. **(B)** Quantification of the number of PC-positive cells after 24h serum starvation in RPMI or DMEM, in both control (blue) and *Ift88*-deficient Col6a1-fibroblasts (red). **(C)** Immunofluorescence analysis showing CF of Col6a1cre^Tg/0^ *Ift88* ^+/+^ Rosa ^mT/mG^ and *Ift88*-deficient *Col6a1cre^Tg/0^Ift88^flx/flx^Rosa^mT/mG^* mice, immunostained for GFP (green), for Arl13b to identify PC (red) and *Ift88* (cyan). *Ift88*-positive PC are indicated with arrows and *Ift88*-negative PC are indicated with asterisks. Bar represents 10µM. **(D)** Schematic representative of the pyruvate treatment timeline in mice. After weaning, control and *Ift88*-deficient mice were treated for 8 weeks with either 0.2M of sodium pyruvate (NaPyr) or sodium chloride (NaCl) as control, added to drinking water. **(E)** Quantification of the percentage of PC-positive colonic fibroblasts in colons of control and *Ift88*-deficient mice : NaCl treatment for control (blue) and *Ift88*-deficient mice (red) and NaPyr treatment in purple. Individual data points represent male (squares) or female (circles) mice. Mean values are indicated. **(F)** Sodium pyruvate treatment increased PC length *in vivo*. Quantification of PC length in colons of control mice under NaCl treatment (blue) or NaPyr treatment (purple). Individual data points represent one mouse (n=5 for NaCl and n=6 for NaPyr). Statistical analysis: *p<0.05, **p< 0.01 by two-tailed unpaired t-test.

### Pyruvate attenuates susceptibility to induced colitis in Col6a1cre-Ift88^flx/flx^-ROSA^mT/mG^

We have recently shown that reduced number of PC in collagen6a1-positive colonic fibroblasts correlates with an increased susceptibility to chemically-induced colitis. In this protocol, mice are treated for 7 days with dextran sulfate sodium (DSS) added to the drinking water. DSS is toxic to mucosal epithelial cells specifically in the colon, but not genotoxic, leading to the destruction of the mucosal barrier and inflammation ^18^. To test whether pyruvate treatment can alter the susceptibility to DSS-induced colitis, *Col6a1cre-Ift88^flx/flx^-ROSA^mT/mG^*mice received sodium chloride or sodium pyruvate supplemented in the drinking water for 8 weeks before DSS treatment (Fig. 7A). At the end of the protocol mice were sacrificed and colons collected for histological analysis. Remarkably, sodium pyruvate treated *Col6a1cre-Ift88^flx/flx^-ROSA^mT/mG^* mice displayed reduced inflammatory signs compared with NaCl-treated mice, including less body-weight loss and increased colon length (Fig. 7B-D). Mucosal architecture of pyruvate treated *Col6a1cre-Ift88^flx/flx^-ROSA^mT/mG^* colons revealed decreased aggressive ulceration. Epithelial changes of pyruvate treated *Col6a1cre-Ift88^flx/flx^-ROSA^mT/mG^* colons exhibited less detrimental crypt abscesses and more active regeneration (Fig. 7E). Notably, CFs from mice treated with sodium pyruvate showed significantly higher PC levels than NaCl-treated animals (Fig. 7F). Thus, treatment of *Ift88*-deficient mice with sodium pyruvate prior and during DSS administration reduced their susceptibility to acute colitis by promoting maintenance of PC-positive col6a1-positive fibroblasts in the colon.

**Figure 7:**
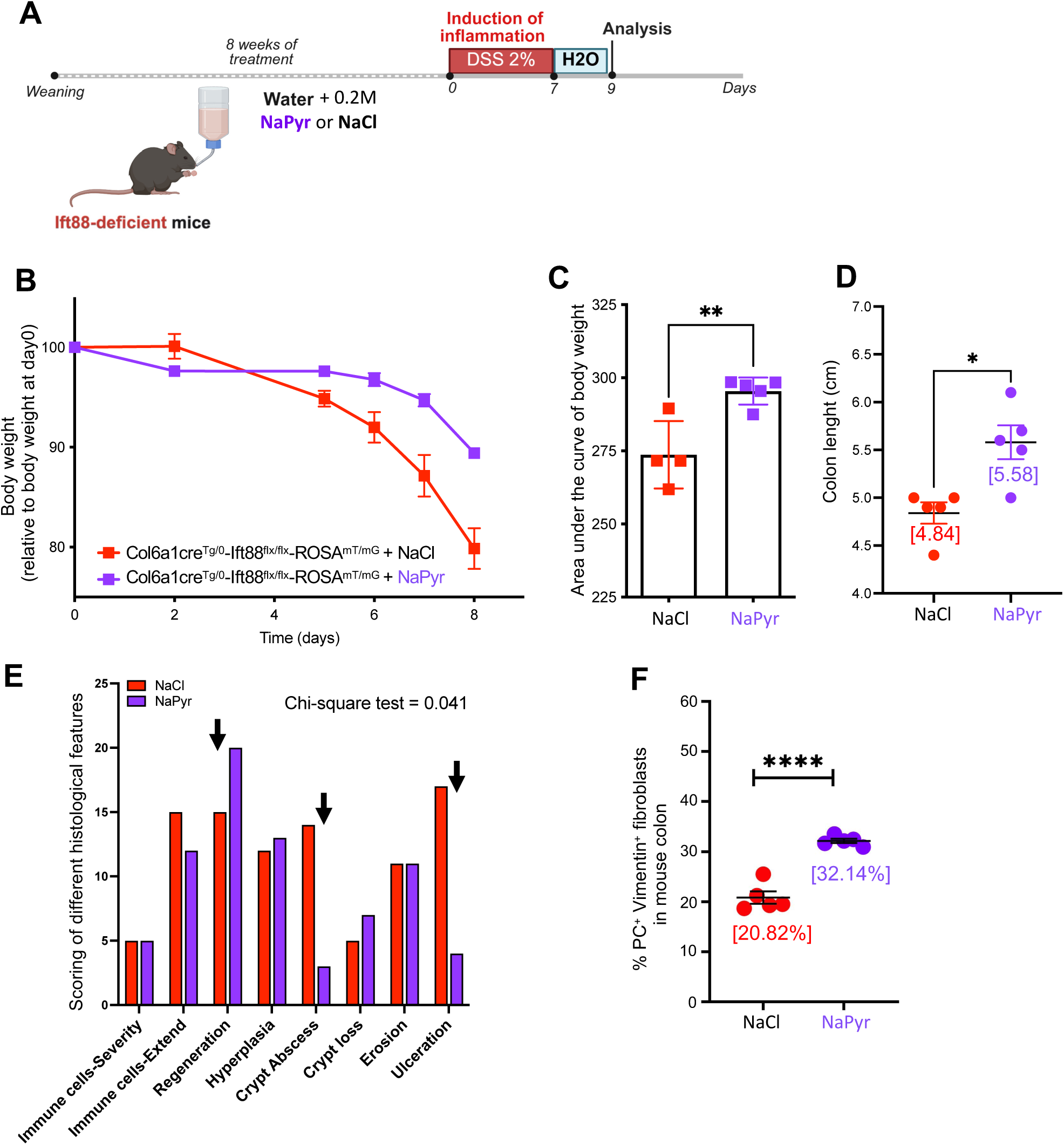
Pyruvate protects Ift88-deficient mice against DSS-induced colitis. **(A)** Schematic representation of the pyruvate treatment timeline in mice exposed to DSS-induced colitis. After weaning, *Ift88*-deficient mice were treated for 8 weeks with either 0.2M of sodium pyruvate (NaPyr) or sodium chloride (NaCl) added to the drinking as well as during the 7-days DSS-treatment. Colons were collected at day 9 after initiation of DSS treatment. **(B)** Body weight curve of *Col6a1cre Ift88*-deficient mice during DSS treatment, showing NaCl treated mice (in red), NaPyr treated mice (in purple). **(C, D)** Graphs displaying respectively the area under the curve of the body weight **(C)** and the colon length **(D)** in treated *Col6a1cre Ift88*-deficient mice. **(E)** Histological scoring of DSS-induced colitis assessing NaCl treated (in red) and NaPyr (in purple) *Ift88*-deficient treated mice. Colon tissue sections were examined and analyzed for histopathological scoring features including immune cells infiltration (severity and extend), regeneration, hyperplasia, crypt abscess, crypt loss, erosion, and ulceration. Arrows highlight the features most impacted by NaPyr treatment. **(F)** Quantification of the percentage of PC-positive colonic fibroblasts in colons of *Ift88*-deficient female mice : NaCl treatment for control (red) and NaPyr treatment (purple). Mean values are indicated. **(C, D, F)** Statistical analysis: two-tailed unpaired t-test; *p<0,05, **p< 0.01, ****p<0.0001.

## Discussion

Here we show that pyruvate is an important regulator of PC biology in CF. We demonstrate that the addition of 2mM pyruvate elongates PC in serum-starved CF-cultures. Importantly, while increasing pyruvate concentrations up to 10 mM did not further extend ciliary length, it significantly increased the proportion of cells that were ciliated.

Previous studies have shown that pyruvate is a major source of acetyl-CoA, which serves as a substrate for the acetylation of both α-tubulin and histones, catalyzed by α-tubulin N-acetyltransferase 1 (ATAT1) and histone acetyltransferases (HATs), respectively. Consistent with this, we found that pyruvate supplementation increased tubulin acetylation in PC and that this modification appears to positively regulate ciliogenesis. This concurs with reports showing that increased tubulin acetylation, catalyzed by ATAT1, can promote cilium assembly ^19,20^. Nevertheless, PC can still form when tubulin acetylation is absent, as demonstrated in ATAT1-deficient mice^21^. Thus, tubulin acetylation is thought to act primarily by protecting cilia from disassembly rather than being strictly required for their assembly. Indeed, a major role for the deacetylase HDAC6 in cilia disassembly has been previously demonstrated ^22,23^. This model aligns with our observation that elevated tubulin acetylation enhances the proportion of ciliated CFs.

Using RNA sequencing we found that pyruvate treatment led to an increase in the expression of MAPK pathway components, a result independently supported by mass spectrometry data. This is consistent with earlier work showing that MAPK signaling modulates PC length in endothelial cells^24^. Proteomics further indicated enhanced TCA cycle flux and electron transport chain activity, indicating increased pyruvate metabolism, as well as enrichment of cilium assembly pathways.

Apart from MAPK pathway, the RNA-sequencing also suggested that pyruvate treatment results in increased histone acetylation, which was validated through the use of specific histone acetylation inhibitors. One possibility is that histone acetylation promotes chromatin accessibility leading to increased transcription, and thus promoting to MAPK pathway activation in CF. However, another study reported that MAPK pathway can activate histone acetylation at gene promoters and thus increase transcription ^25^. Furthermore, a recent study showed that histone acetylation may in turn enhance expression of MAPK pathway components, forming a positive feedback loop ^26^. Additional experiments are required to decipher whether such feedback mechanism operates in CF or whether the effect of pyruvate is more direct leading to increased histone acetylation, which in turn promotes the expression of MAPK pathway components. Nonetheless, our finding that histone acetylation contributes to pyruvate-driven ciliogenesis is in agreement with previous reports indicating that such modification can modulate transcriptional programs involved in PC formation. For example, depletion of the histone acetyltransferase KAT2B (lysine acetyltransferase 2 B) in mouse embryonic fibroblasts impaired ciliogenesis ^27^. This concurs with additional reports linking histone acetylation with cilia formation ^28,29^.

Intriguingly, we observed that inhibition of the histone acetyltransferase EP300 reduced pyruvate-induced PC elongation to a similar extent as MAPK inhibition, and decreased pyruvate-stimulated ciliogenesis in a manner resembling ATAT1 inhibition. These findings imply that EP300 acts upstream, or in coordination with, the MAPK pathway and ATAT1 to regulate pyruvate-dependent ciliogenesis and cilium elongation.

Apart from acetylation we also found that another α-tubulin post-translational modification called detyrosination played a role in the ciliary length regulation in PC stability ^30,31^. Detyrosination is primarily catalyzed by enzymes belonging to vasohibin family, VASH1 and VASH2 ^15,16^, and previous studies have linked the accumulation of this modification with PC elongation ^32,33^. For example, MDCK cells genetically modified to inactivate the Tubulin Tyrosine Ligase (TTL) enzyme, which reverses detyrosination by adding a tyrosine residue, displayed elevated levels of detyrosination and assembled longer PC. A more recent study in the green alga Chlamydomonas reinhardtii showed that Vash deletion affects anterograde IFT train movement, leading to ciliary elongation ^33^. Consistent with these findings, we observed that treatment of ciliated CF cells with LV80, a potent VASH inhibitor^34^ did not alter the proportion of ciliated cells but did negatively affect PC length. Interestingly, while the addition of LV80 reduced PC length under both 2 mM and 10 mM pyruvate conditions, a significant decrease in the levels of tubulin detyrosination was observed only at 2mM pyruvate. This apparent discrepancy can be explained by the higher sensitivity of detecting changes in PC length compared with bulk detyrosination levels. In support of this notion, the relative change in PC length caused by LV80 was more pronounced at 2mM pyruvate than at 10mM (0.6-fold vs. 0.9-fold, respectively).

Similarly, MEK and ERK1/2 inhibition decreased PC detyrosination and reduced ciliary length. In contrast, inhibition of tubulin acetylation affected number of cilia without influencing detyrosination status or ciliary length. Taken together, these findings support a model in which pyruvate regulates PC length primarily through MAPK-dependent changes in tubulin detyrosination at the cilium, whereas tubulin acetylation contributes to ciliogenesis but not elongation.

Finally, we show that pyruvate treatment rescues PC deficiency in *Ift88*-depleted fibroblasts *in vitro* and *in vivo*. This result is particularly striking considering that IFT88 is thought to be essential for ciliary assembly in various model organisms ^17^. Previous work has suggested that ciliation can be partially restored in the absence of IFT88 through pharmacological intervention. Spasic *et al.* showed that fenoldopam, a dopamine D1 receptor agonist, restored both the presence and the length of cilia in IFT88-silenced osteocytes without increasing IFT88 expression ^35^. This report supports our finding that pyruvate supplementation can restore ciliogenesis in *IFT88*-deficient fibroblasts. Future studies will reveal whether the capacity of pyruvate to overcome IFT88 dependency is specific to collagen VI–positive fibroblasts or also applies to other cell types. Indeed, a recent study demonstrated that a substantial fraction of the ciliary proteome is cell type–specific ^36^. It is therefore conceivable that the capacity of pyruvate supplementation to compensate for IFT88 deficiency is restricted to specific cell types, such as CF.

Remarkably, in mice lacking *Ift88* specifically in collagen 6a1–positive fibroblasts, pyruvate treatment restored primary cilia and mitigated DSS-induced colitis. These results are consistent with our previous report that PC loss in collagen 6a1–positive fibroblasts, caused by either Ift88- or Kif3a-deficiency, promotes colonic inflammation ^6^. Together, these observations support a model in which primary cilia act as regulators of intestinal inflammation. Nevertheless, we do not exclude the possibility that pyruvate may modulate susceptibility to DSS-induced colitis through additional, PC-independent mechanisms. Given its capacity to restore ciliation and ameliorate inflammatory outcomes, pyruvate supplementation may hold therapeutic potential not only in colon inflammation but also in other cilia-related disorders.

## Acknowledgments

We are grateful to the excellent platforms in Montpellier: the “Réseau d’Histologie Expérimentale de Montpellier” - RHEM facility supported by SIRIC Montpellier Cancer (Grant INCa_Inserm_DGOS_12553), the European regional development foundation and the occitanian region (FEDER-FSE 2014-2020 Languedoc Roussillon) for processing our animal tissues, histology technics and expertise ; the imaging facility MRI, member of the national infrastructure France-BioImaging (https://ror.org/01y7vt929) supported by the French National Research Agency (ANR-24-INBS-0005 FBI BIOGEN)" ; the animal experimentation platform “Réseau des Animaleries de Montpellier” - RAM, as well as the IGMM mouse facility “Zone d’Expérimentation et de Formation de l’IGMM” - ZEFI. Many thanks to Thierry Gostan and the platform SERANAD for help with the statistical analysis and the support of the funding organizations Cancer Inserm-ITMO Aviesan 2021 (to MH, EM), ANR (AAPG2021, CILCOL to MH, EM), and FRM (“équipe labellisé”). The authors are thankful to Professors Kollias (BSRC Fleming Greece) and Prof. Yoder (The University of Alabama at Birmingham), for providing Col6-Cre and IFT88flx/flx strains, respectively. Computing resources for the RNA-seq primary analysis were provided by the computing facilities DISC (Datacenter IT and Scientific Computing) of the Centre de Recherche en Cancérologie de Marseille.

## Methods

### Animals

Mouse experiments were performed in strict accordance with the guidelines of the European Community (directive n°2010/63/EU) and the French National Committee (Project number APAFIS#18685 and #52137) for the care and use of laboratory animals and were approved by the Regional Ethics committee. Statistical analysis was conducted to minimize the number of mice to be used for identifying significant differences. To study PC, we used *Ift88^flx/flx^ Col6a1cre* mice deficient for PC in collagen 6a1 positive fibroblasts, as previously described. To track this fibroblast subset, we crossed the *Ift88^flx/flx^ Col6a1cre* mice with mice expressing *Rosa mT/mG* as tracker. The *Rosa^mT/mG^* mice ubiquitously express a red fluorescent protein (Tomato) before any recombination. In cells where Cre recombinase is active, the Tomato cassette is excised and a membrane-targeted green fluorescent protein (GFP) is expressed instead, allowing robust visualization of knockout cells and their lineages in tissue by immunofluorescence. *Ift88^flx/flx^ Col6a1cre Rosa^mT/mG^*mice were maintained on mixed 129 and C57BL/6 genetic background, and experimental groups including littermates were housed together according to gender and age. Genotyping was done using primers listed in Table 1. Mice used for the *in vivo* experiments were used at age of 8-12 weeks.

**Table 1:**
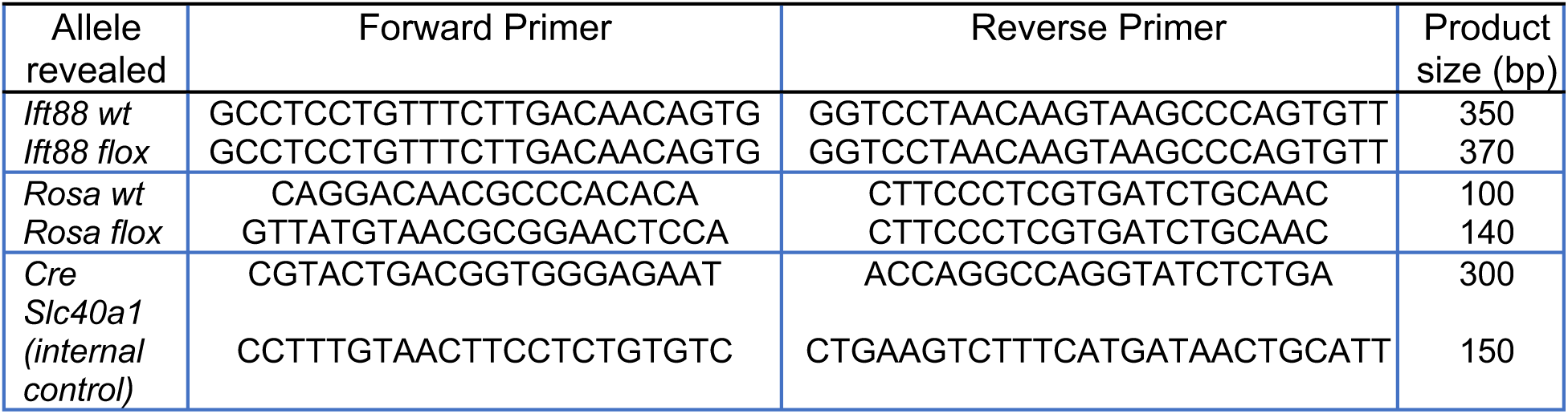
Sequences of the primers used for mouse genotyping.

### Reagents

The MEK1/2 inhibitor PD98059 (used at 20µM), ERK1/2 inhibitor SCH772984 (used at 300nM), the inhibitor of pyruvate transport at the plasma membrane AZD3965 (used at 200nM), the microtubule acetylation inhibitor GM-90257 (used at 500nM) and the histone acetyl transferase inhibitors for E1A-associated Protein p300 (Ep300) C646 (used at 25µM) were purchased from MCE (MedChem Express, Monmouth Junction, NJ, USA). The Vash inhibitor LV80 was used at 100µM and has been previously described ^37^. The inhibitor of the Mitochondrial Pyruvate Carrier (MPC), UK-5099 (used at 50 µM) and sodium acetate were purchased from Sigma-Aldrich (UK-5099: PZ0160; sodium acetate: S2889). Sodium pyruvate (Gibco, 11360070) was used for cell culture, and sodium pyruvate (Sigma-Aldrich, P2256) was used for *in vivo* treatment. These compounds were dissolved in Dimethyl Sulfoxide (DMSO) and kept frozen at -20°C as stock solutions. Working solution were made as dilution of the stock solutions with medium and the final DMSO concentration in the medium did not exceed 0.2% (v/v). At this concentration, DMSO itself had no effect on cell viability, cilia length, number or morphology of PC-positive CF in the respective assays (Fig. 4).

### Colonic primary fibroblast isolation and culture

The protocol for isolating colonic fibroblasts (CF) was modified according to ^6^. Briefly, colons were isolated from mice (38-42 days old male mice), flushed with Phosphate Buffered Saline (PBS) to remove feces, and then opened longitudinally. Colons were then cut into small pieces, extensively washed, and incubated in Ethylene Diamine Tetra-acetic Acid (EDTA)-containing buffer Hank’s Balanced Salt Solution (HBSS (ThermoFisher,14025050), 2% Fetal Calf Serum (FCS), 5 mM EDTA) at 37°C for 20 minutes with continuous shaking to release colonic epithelial cells. After extensive washing, the tissue was further minced into smaller pieces and digested enzymatically in HBSS - 2% FCS solution containing 62.5 µg/ml Liberase (Sigma, 05401020001) and 40 µg/ml DNase (Sigma, 11284932001). Digestion was performed at 37°C for 50 minutes under continuous shaking with a magnetic stirring device to isolate CF. Digestion was stopped by adding Dulbecco’s Modified Eagle Medium (DMEM) medium supplemented with 10% FCS. Cells were centrifuged at 300g for 7 minutes at 4°C. The pellet was resuspended with PBS and filtered through a 100µm strainer, to remove undigested pieces. The filtered cell suspension was centrifuged again at 300g for 7 minutes at 4°C. For the culture, cells were filtered through a 40µm strainer to obtain single cell suspension, and approximately 3 x 10^6^ cells were seeded per well in a flat-bottom 24-well plate. Cells were cultured in DMEM (250mg/ml Strepto-7.5×10^6^U/ml PenicillineG peni/stepto, 10%FCS (Sigma, BCBZ3745), 10mM 4-(2-HydroxyEthyl)-1-PiperazineEthaneSulfonic acid (HEPES) (Gibco, 15630080), 2mM sodium pyruvate, 50µM beta-mercaptoethanol (Gibco, 31350010)) for 2 passages and maintained at 37°C in a humidified atmosphere with 5% CO2. The culture medium was replaced every 2 days. The first passage was performed between days 5 and 7 of culture, and the second passage between days 9 and 11.

### Serum Starvation and chemical treatments

At passage 2, CF were seeded on 2% gelatin-coated glass coverslip (Sigma, F8775) in a flat-bottom 24-well plate. Around 14-17 days of culture, when CF reached a confluence of 70–90%, the culture medium was changed for and additional 24 hours into serum-free DMEM or Roswell Park Memorial Institute (RPMI) medium containing 1% Bovine Serum Albumin (BSA) fraction V (Euromedex, 04100812E) and 50µM β-mercaptoethanol (Gibco, 31350010) to induce PC formation. During the serum starvation treatment, cells were treated with sodium pyruvate or sodium acetate (0, 2, 5 or 10mM), as well as the different inhibitors described previously.

### Immunofluorescence on cells

At the end of treatment, cells present on coverslips were washed once with PBS, and fixed with 4% Paraformaldehyde (PFA; Sigma G9391) for 10 minutes at room temperature (RT). After fixation, cells were permeabilized and blocked in PBS containing 3% BSA (Sigma, 0000383235) and 0.3% Triton X-100 (Sigma, 61K0173) for 30 minutes at RT. The same buffer (PBS - 3% BSA - 0.3% TritonX-100) was used for washes as well as for primary and secondary antibody dilutions. Cells were incubated with primary antibodies for 2 hours at RT, washed five times, and then incubated with fluorescent-labeled secondary antibodies for 1 hour at RT in the dark. Nuclei were stained with 10 µg/ml of 4’,6’-diamidino-2-phenylindole (DAPI, Sigma, MBD0015) for 5 minutes, followed by four washes. Coverslips were mounted using ProLong™ Gold Antifade Mountant (Invitrogen, 11539306). The list of antibodies used is described in Table 2.

**Table 2:** Primary and secondary antibodies used in Immunofluorescence staining.

| Protein name | Supplier | Reference | Host | Dilution |
| --- | --- | --- | --- | --- |
| <u>Primary antibodies</u> |  |  |  |  |
| *Arl13b* | Neuromab | N295B/66 | Mouse | 1/1000 |
| *Arl13b* | Proteintech | 17711-1-AP | Rabbit | 1/1000 |
| *Acetyl tubulin* | Sigma | T6793 | Mouse | 1/1000 |
| *IFT88* | Proteintech | 13967-1-AP | Rabbit | 1/100 |
| GFP | Novus Biologicals | NB100-1770 | Goat | 1/500 |
| *De-tyrosinated alpha Tubulin* | Invitrogen | RM444 | Rabbit | 1/500 |
| *Vimentin* | BioLegend | Poly29191 | Chicken | 1/1000 |
| *Vimentin* | Cell Signaling | D21H3 | Rabbit | 1/200 |
| <u>Secondary antibodies</u> |  |  |  |  |
| Anti-rabbit Alexa A488 | Invitrogen | A21206 | Rabbit | 1/1000 |
| Anti-goat Alexa A488 | Invitrogen | A11055 | Goat | 1/1000 |
| Anti-mouse Alexa A488 | Invitrogen | A11017 | Mouse | 1/1000 |
| Anti-chicken Alexa A488 | Invitrogen | A78948 | Chicken | 1/1000 |
| Anti-mouse Alexa A555 | Invitrogen | A31570 | Mouse | 1/1000 |
| Anti-rabbit Alexa A647 | Invitrogen | A31573 | Rabbit | 1/1000 |

Images were acquired using a Zeiss Laser Scanning Microscopy (LSM) 980 confocal microscope equipped with AiryScan 8Y detection and a 40X oil immersion objective. Eight consecutive Z-stack sections were captured at 0.29 µm intervals over a 2.03 µm range. Image reconstruction was performed using Zeiss Zen software. Image analysis was done on at least five random fields (visualizing around 100 cells/field) per sample using Fiji (Image J, NIH). PC were manually counted, and their length was measured by dragging a line with a segmented-line tool along each Arl13b stained-cilium and measuring the line length. PTM intensity was quantified in PC by performing integrated density over PC area. The area of the PC was delineated using Arl13b staining and the density of PTM staining was quantified in this area (Supplementary Fig. 9).

### NaPyr/NaCl treatment of the mice and DSS induced acute colitis

Immediately after weaning, mice were divided into two experimental groups. The treatment group receiving 0.2M of sodium pyruvate while the control group received 0.2M of sodium chloride, both diluted in drinking water to have an osmolarity match control. Mice were treated for 8 consecutive weeks and solutions were freshly prepared and changed every two days. Body weight was measured regularly during this treatment period to monitor the effect of sodium pyruvate in healthy mice. Following the eight weeks of pyruvate treatment, at around 11-12 weeks of age, mice were treated for 7 days with 2% (w/v) dextran sodium sulfate (DSS, TdB Sweden) added to the drinking solution, which continued to contain either sodium pyruvate or sodium chloride. Body weight was recorded daily during the all experiment, and in case weight loss was more important than 20%, mice were immediately sacrificed. On day 8, mice received normal drinking water, they were sacrificed on day 9 and colons were collected. Body weight curves for both the pre-treatment phase and the DSS-treatment phase were generated. For the analysis of DSS-induced acute colitis, disease severity was assessed by calculating the Area Under the Curve (AUC) of body weight change, and by measuring colon length and spleen weight.

### Tissue collection and paraffin embedding

At the end of *in vivo* protocols, colons were collected, flushed with PBS, rolled up flat and embedded in paraffin after fixation in 10% neutral buffer formalin for 24 hours at room temperature. Colons sections of 4µm were obtained using microtome Niagara and mounted onto glass microscope slides. Slides were used for immunofluorescence analysis or for histological examination after Hematoxylin-Eosin-Saffron (HES) staining.

### HES and Histological Scoring

HES staining was done by the platform Réseau d’Histologie Expérimentale de Montpellier (RHEM). Slides were scanned using the Nanozoomer scanner (Hamamatsu Photonics, Hamamatsu City, Japan) and analyzed with Nanozoomer Digital Pathology (NDP) Viewer. HES-stained slides were evaluated based on histological scoring system adapted for DSS-induced acute colitis described by *Erben et al.* ^38^. Histological assessment of DSS-impact on colon considers three categories reflecting the severity of the induced colitis: inflammatory cell infiltration, epithelial changes and mucosal architecture changes as described in Supplementary Table 2.

### Immunofluorescence on paraffin tissue section and image analysis

Colons slides were deparaffinized in xylene (10 minutes), then gradually rehydrated in 100% ethanol (5 minutes), 95% ethanol (3 minutes), 70% ethanol (3 minutes) and H2O (5 minutes). After incubation in EDTA 1mM pH9.0 for 30 min at 99°C (antigen retrieval), slides were washed twice in PBS. Tissues were blocked in PBS - 5% BSA - 0.3% TritonX-100 for 1 hour in humid chamber at RT. The same buffer (PBS - 5% BSA - 0.3% TritonX-100) was used for primary and secondary antibody dilutions. Tissues were incubated with primary antibodies overnight 4°C, washed three times in PBS, and then incubated with fluorescent-labeled secondary antibodies mixed to DAPI (10 µg/ml) for 1 hour at RT in the dark in humid chamber. After four washes, slides were mounted with cover glasses using Polong^TM^ Gold Antifade Mountant. The list of antibodies used is described in Table 2.

Images were acquired using an Andor Dragonfly Spinning Disk Confocal inverted microscope using a 40X oil immersion objective. 26 Z-stack sections were captured at 0.2µm intervals over a 5µm range. Quantification of PC number in vimentin-positive fibroblasts was done on at least five random fields (about 1 mm^2^ each) per sample and were analyzed using Fiji software. Quantification of the length of Arl13b positive - PC was performed using Fiji software and special plugins Labkit and MorpholiJ. Images were first segmented using the Labkit plugin, which enables interactive machine learning-based pixel classification to accurately delineate PC within the image. Segmentation models were trained on representative regions to distinguish PC from background and to take in account only PC in vimentin positive fibroblasts. Following segmentation, quantitative length measurement of PC was performed using MorpholibJ plugin using ellipsoid features. Using the major axis of the equivalent ellipsoid fitted to each PC, as computed by MorphoLibJ’s region analysis tools, the length of each PC was accurately estimated as twice the largest ellipsoid radius (Length=2×Rmax).

### RNA extraction

The Roche® high pure RNA isolation kit was used for RNA extraction. Cell pellets were resuspended in 200μl PBS then 400μl lysis buffer. The suspension was vortexed 20 seconds, transferred onto a spin column coupled to a collection tube and centrifuged at 8000g for 30 sec. After discarding the flow through, 100μl DNAse I was added to each spin column and incubated at RT for 15 minutes for genomic DNA removal. The columns were then centrifuged as described and the flow through discarded. Columns were washed as indicated by the supplier and RNA was eluted into Eppendorf® DNA-LoBind tubes using 50μl elution buffer and centrifuged at 8000g for 1 min. RNA concentration was determined using Nanodrop before to be used for RNA sequencing.

### RNA Sequencing using Dr TOM software

Library preparation and sequencing were performed by Beijing Genomics Institute (BGI) (Shenzhen, China). Differential gene expression analysis was followed by functional enrichment analysis, Kyoto Encyclopedia of Genes and Genomes (KEGG) pathway (https://www.kegg.jp/) enrichment analysis of annotated different expressed genes was performed by Dr. Tom platform (https://biosys.bgi.com/). The significant levels of terms and pathways were corrected by Q value with a rigorous threshold (Q value < 0.05).

Gene Set Enrichment Analysis (GSEA) results were displayed as enrichment plots showing the ES curve, with Normalized Enrichment Score (NES) values used to annotate and quantify the normalized enrichment level. The leading-edge subset of genes was also identified to highlight core contributors to the enrichment signal.

The RNA-seq data generated in this study have been deposited in NCBI’s Gene Expression Omnibus and are accessible through GEO Series accession number GSE311989.

### Shotgun label-free proteomics data acquisition and processing

In label-free shotgun proteomic analysis, samples were lysed by ultrasonication (80% amplitude, Sonicator U200S control, IKA Labortechnik, Staufen, Germany) in lysis buffer (urea 6 M in 100 mM Tris/HCl, p H = 7,5) and 25 µg of protein for each condition were digested by trypsin (0,5 µg/µl Trypsin, Sigma-Aldrich, St. Louis, MI, USA) according to the single-pot, solid-phase-enhanced sample-preparation (SP3) protocol, previously described ^39^ and (collaborator’s manuscript submitted in September - Acta Neuropathologica Communications). Proteomics data were acquired according to Di Stefano *et a.l* ^40^ by using the UltiMateTM 3000 UPLC (Thermo Fisher Scientific, Waltham, MA, USA) chromatographic system coupled to the Orbitrap FusionTM TribridTM (Thermo Fisher Scientific) mass spectrometer. Proteomics raw data were processed using Thermo Proteome Discoverer (PD) version 2.4.0.305 (Thermo Fischer Scientific) to characterize the proteome composition of Sample 1 and Sample 2 and to identify differentially expressed proteins between them, as described in Ricciardi *et al.* ^41^. Proteins dataset, containing the protein ID and the expression ratio, was uploaded to the Ingenuity Pathway Analysis tool (IPA, Qiagen, Hilden, Germany) to perform a gene ontology and functional enrichment analysis based on all quantified proteins. Raw proteomics data are available via the ProteomeXchange Consortium with the dataset identifier PXD071337.

### Statistical analysis

All statistical analysis of quantifications for the mouse study were performed with GraphPad Prism version 10 using an unpaired student t-test (2-tailed) when data passed the normality tests (Kolmogorov-Smirnov (KS) normality test, D’Agostino–Pearson omnibus normality test, Shapiro–Wilk normality test). If values were not normally distributed, a nonparametric Mann-Whitney U test was used. Chi-square test (χ2) was used for the contingency of histological scoring (Supplementary table 2).

