## supplemental data for "Pyruvate promotes ciliogenesis bypassing *IFT88* dependency and attenuates DSS-induced colitis"

#### Supplementary information.

Supplementary Figures 1-9 with legends

Supplementary Tables 1 and 2

### Supplementary Figure 1

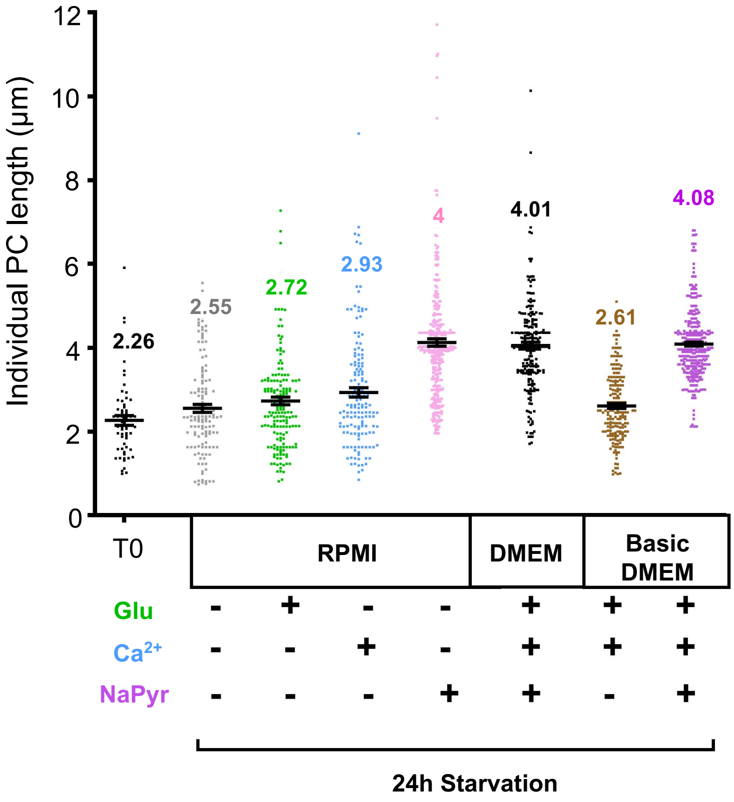


**Supplementary Figure *1*: *Measurement of primary cilium length in CF under varying culture conditions*.** Quantification of PC length on colonic fibroblasts after 24h starvation in RPMI, RPMI supplemented with either Glucose (Glu, 0.14mM), Ca2+ ions (0.93 mM), sodium pyruvate (2mM), or with DMEM or basic DMEM. Each dot represents the length of an individual PC and all dots group together data from 3 different mice.

### Supplementary Figure 2

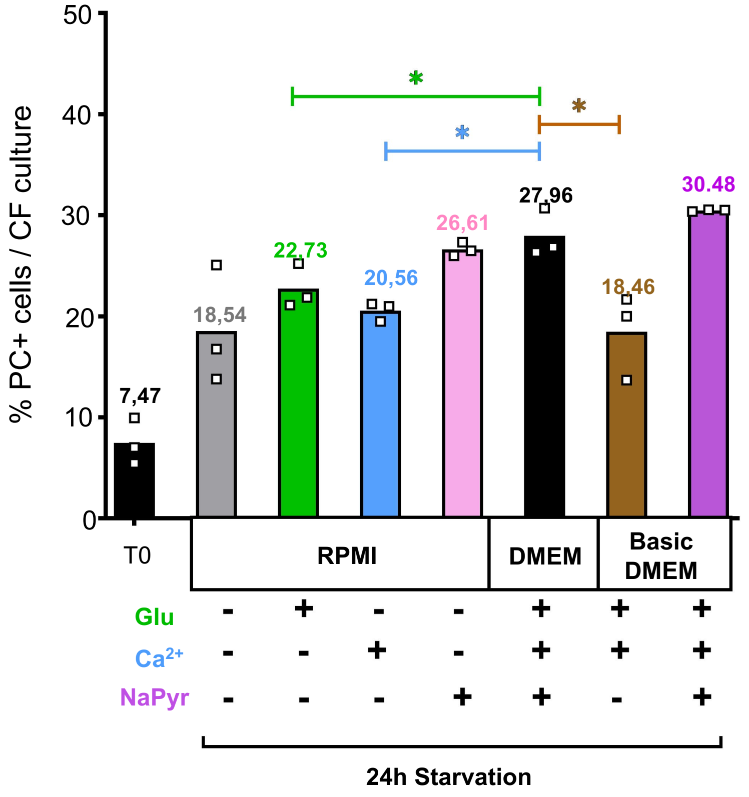


**Supplementary Figure 2**: ***Medium composition reveals pyruvate as a key regulator of PC abundance.*** Quantitation of the percentage of PC-positive cells in CF cultures after 24h starvation in RPMI, RPMI supplemented with either Glucose (Glu, 0,14mM), Ca2 -positive-ions (0,93 mM), sodium pyruvate (2mM), or with DMEM or basic DMEM. Statistical analysis: two-tailed unpaired t-test; *p< 0.05. Mean of 3 independent CF culture are presented per condition.

### Supplementary Figure 3

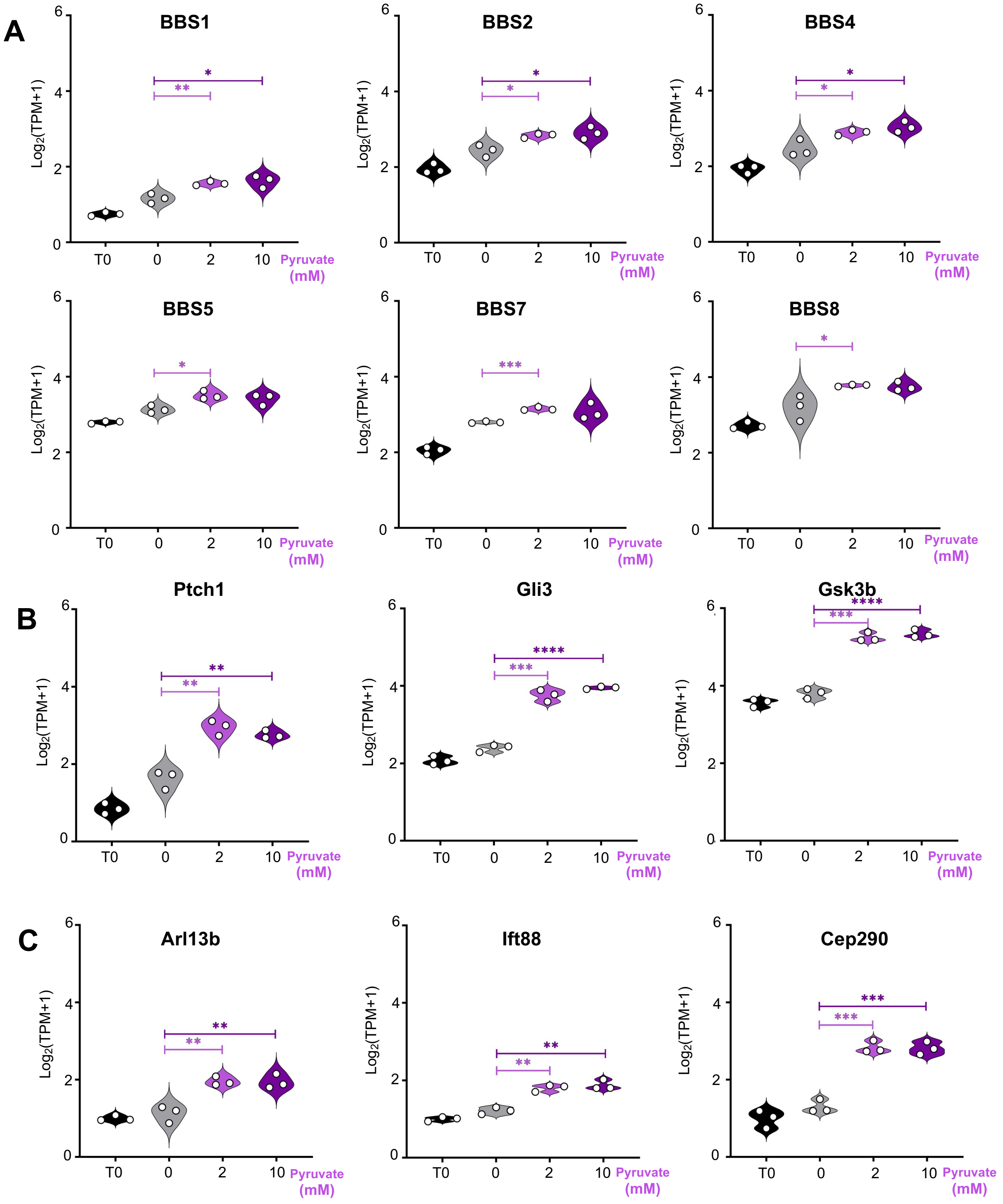


**Supplementary Figure 3:** ***Elevated transcript levels of cilia-related genes in serum-starved colonic fibroblasts in response to pyruvate.*** Transcript levels in serum-starved colonic fibroblasts treated with the indicated concentrations of pyruvate for: BBSome genes **(A)**, hedgehog signaling pathway genes **(B)** and additional primary cilia-associated genes **(C)**.

### Supplementary Figure 4

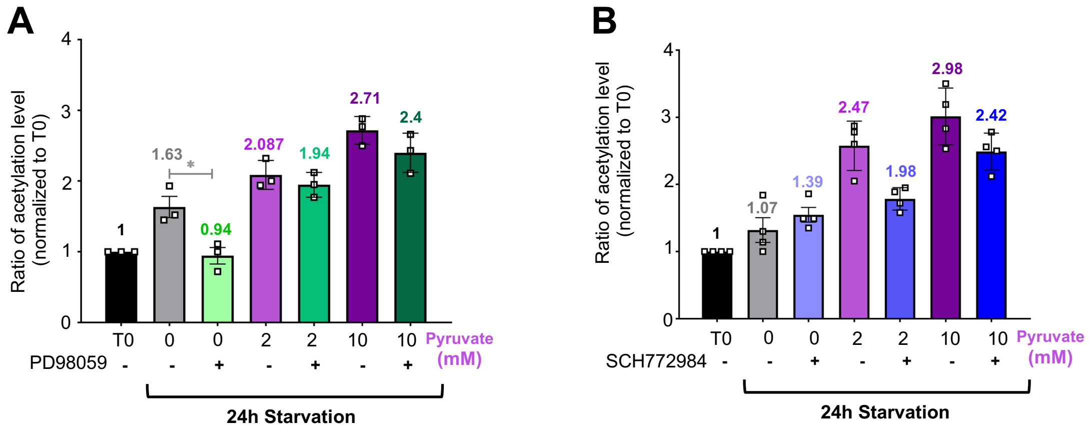


**Supplementary Figure 4:** ***Inhibition of MAPK signaling pathway had no effect on PC acetylation.*** Acetylation levels of PC in 24h serum-starved colonic fibroblasts in absence or in presence of sodium pyruvate and treated with the MEK1/2 inhibitor PD98059 (A) or the ERK1/2 inhibitor SCH772984 (B).

### Supplementary Figure 5

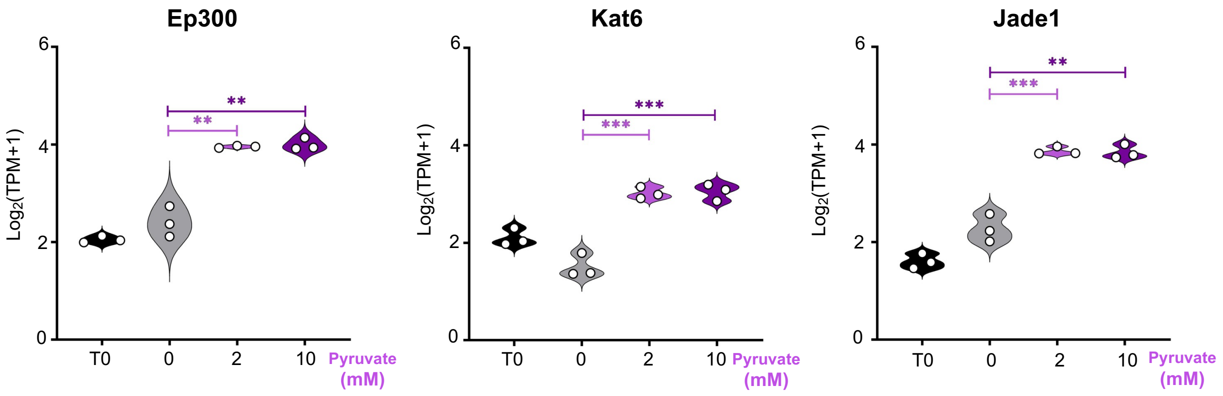


**Supplementary Figure 5:** ***Transcript levels of histone acetyltransferases in serum-starved colonic fibroblasts in the presence of the indicated concentrations of pyruvate.*** RNA-seq analysis shows expression of *Ep300* (left panel), lysine acetyltransferase *Kat6* (middle panel), and the scaffold protein *Jade1*, a component of the histone acetyltransferase complex (right panel). Expression levels are presented as $\log_2$-transformed normalized transcript abundance (Transcripts Per Million, TPM) for improved visualization.

### Supplementary Figure 6

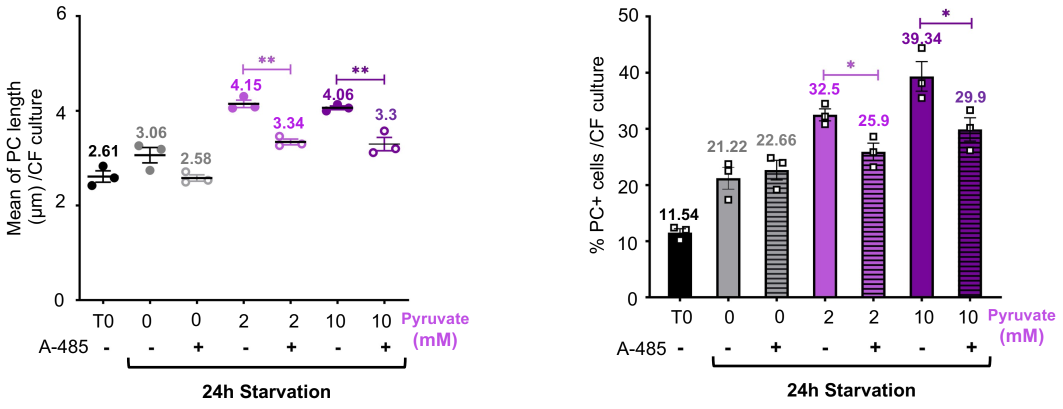


**Supplementary Figure 6: Measurement of Primary cilia length and number in colonic fibroblasts treated with Histone acetylation inhibitor A-485.** *Effect of histone acetylation blocking using the inhibitor A-485 (500nM) on both PC length (left panel) and number of PC-positive cells (right panel). Statistical analysis: two-tailed unpaired t-test; *p<0.05, **p< 0.01.*

### Supplementary Figure 7

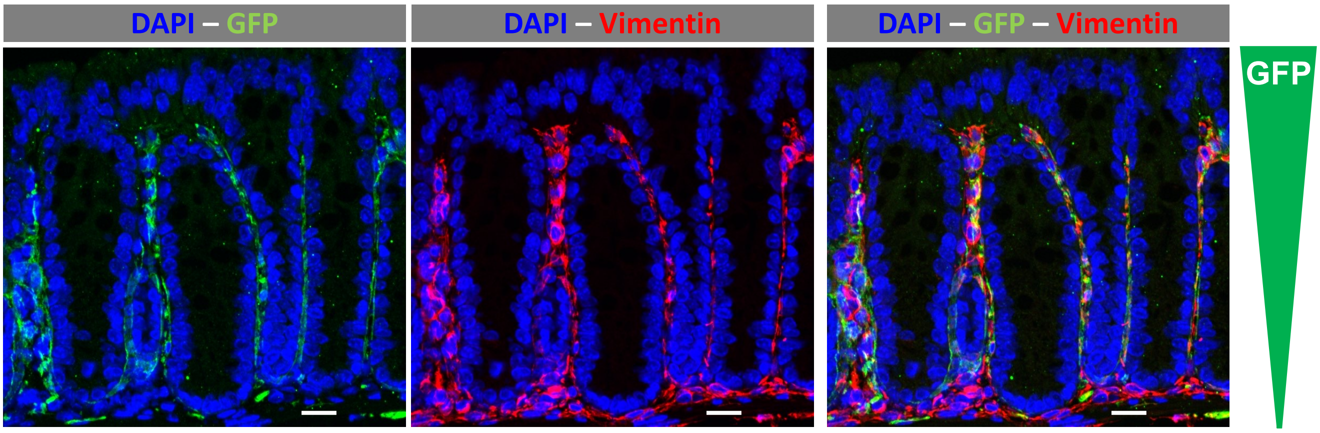


**Supplementary Figure 7:** ***Spatial gradient of GFP-positive Col6a1 fibroblasts in mouse colon.*** Immunofluorescence analysis of Col6a1cre-Ift88$^{flox/flox}$-ROSA$^{mT/mG}$ mouse colon showing GFP-positive Col6a1 fibroblasts enriched in the upper and middle part of the lamina propria surrounding the colonic crypts. Col6a1-positive fibroblasts are marked with GFP (green) and vimentin (red). Scale bar: 20μm

### Supplementary Figure 8

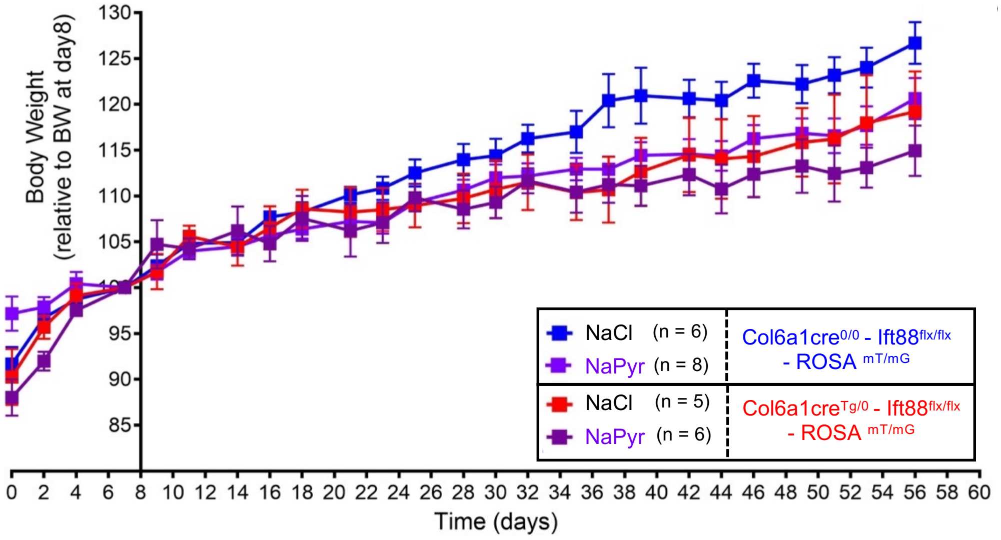


**Supplementary Figure 8 :** ***Body weight curve of control or Ift88-deficient mice treated 8 weeks with sodium pyruvate or sodium chloride solution.*** Graph shows body weight curves of control and Ift88-deficient mice treated with sodium chloride (NaCl, blue and red) or sodium pyruvate (NaPyr, light and dark purple) as indicated. Day0 of the treatment corresponds to 3week of age for the mice, since the treatments started just after weaning. Data shown are representative of three independent experiments. Number of mice per group are indicated.

### Supplementary Figure 9

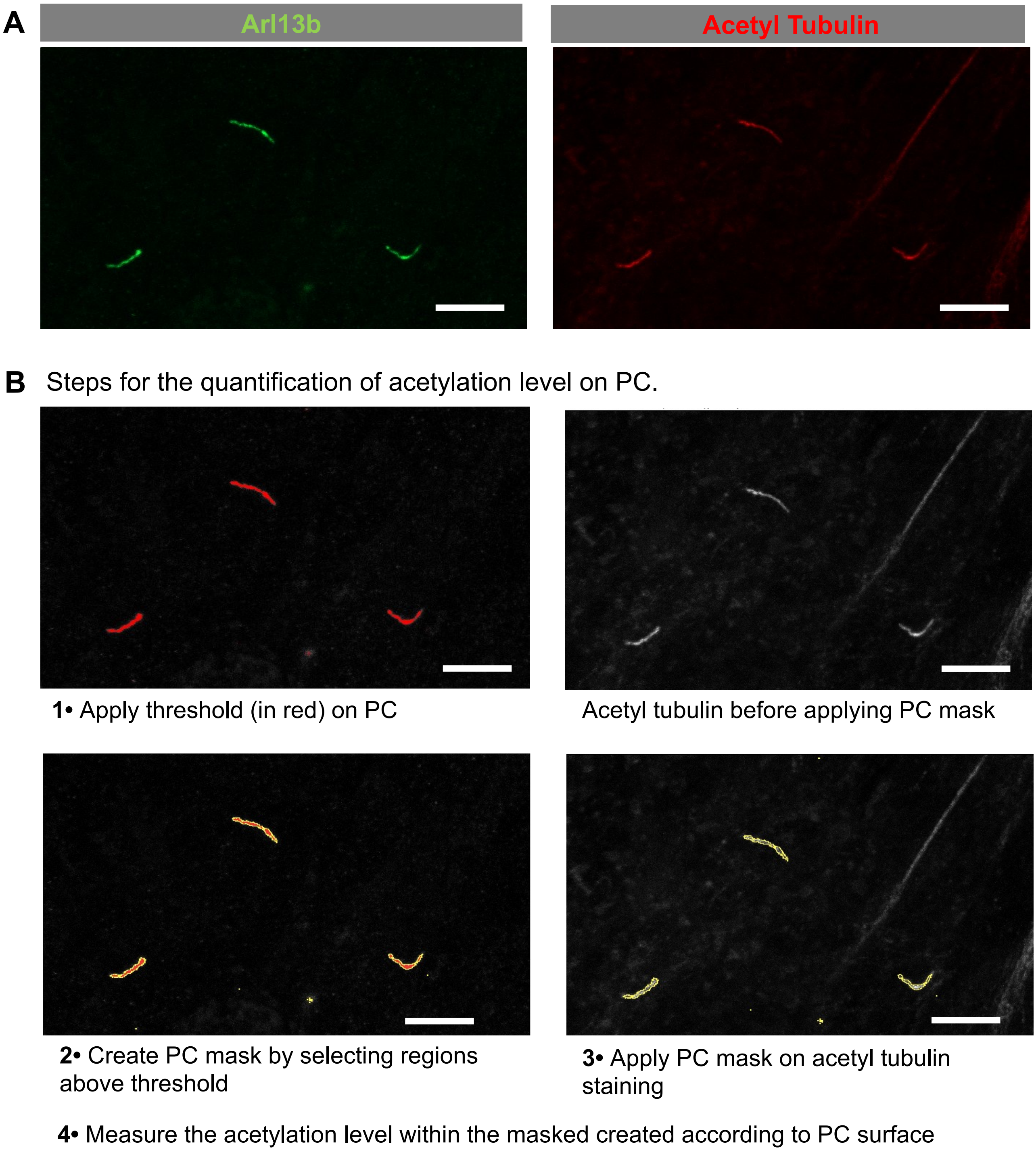


**Supplementary Figure 9:** ***Steps used to quantify acetylation level in PC***
**(A)** Immunofluorescence images used for the quantification: PC in green (Arl13b marker) and acetylation in red (acetyl tubulin antibody). **(B)** Different steps used to quantify acetylation levels of PC using Image J/fiji. **1)** The starting point is the staining of PC by Arl13b, on which a threshold is applied to delineate PC surface as indicated by the red coloration. **2)** A binary PC mask (yellow line) is generated by selecting all pixels above the threshold. **3)** This mask is applied to the acetylated tubulin channel to specifically delineate the acetylated tubulin signal associated with PC. **4)** Acetylation intensity is measured exclusively within the masked PC regions. Scale bars: 5 µm

**Supplementary Table 1:** ***Mass spectrometry analysis revealed TCA cycle, MAPK signaling and cilium assembly among the upregulated pathways***.

Proteomic analysis was performed on 24h serum-starved CF in absence or presence of 2mM pyruvate. Pathways mentioned are upregulated in the pyruvate condition. Sample 1 and 2 were obtained from two distinct CF cultures derived from two individual mice and presented in the table classed by rank and p-value (indicated with -log values).

| Upregulated Pathways | TCA cycle and respiratory electron transport | | MAPK signaling | | Cilium Assembly | |
|---|---|---|---|---|---|---|
| | Rank | -log(p-value) | Rank | -log(p-value) | Rank | -log(p-value) |
| Sample 1 | 8 | 12.5 | 17 | 12.5 | 69 | 5.72 |
| Sample 2 | 5 | 15.3 | 6 | 16.4 | 17 | 5.66 |

**Supplementary table 2: Histomorphological features used for DSS-induced colitis scoring in mouse model.** Table showing the categories, the criteria, their definitions and the associated scores (in parenthesis) calculated from the percentage of each criteria relative to the analyzed surface.

| Category | Criteria | Definition | % of analyzed aera and scoring values |
|---|---|---|---|
| Inflammatory cell infiltration | Severity | Area containing immune cells | <10% (1)<br>>10% (2) |
| | Extend | Expansion of immune cell infiltration | Mucosal (1)<br>Submocosal (2)<br>Transmural (3) |
| Epithelial changes | Hyperplasia | Increase in epithelial cell numbers in longitudinal crypts (including goblet cells); visible as crypt elongation | <10% (1)<br>10-20% (2)<br>20-30% (3)<br>30-40% (4)<br>>40% (5) |
| | Regeneration | Crypts with reduction of goblet cell number that display clear droplets | <20% (3)<br>20-40% (4)<br>>40% (5) |
| | Cryptitis | Immune cells between epithelial cells | <5% (2)<br>>5% (3) |
| | Crypt Abscess | Immune cells in crypt lumen | <10% (3)<br>>10% (4) |
| Musocal Architecture | Crypt loss | Mucosa devoid of crypts | <10% (1)<br>10-20% (2)<br>>20% (3) |
| | Erosion | Mucosa without crypts or surface epithelium | <10% (2)<br>10-20% (3)<br>>20% (4) |
| | Ulceration | Mucosa without crypts or surface epithelium associated with broken muscularis mucosa and/or hemorrhage | <10% (3)<br>10-20% (4)<br>>20% (5) |
